# Temperature outweighs diet in shaping developmental performance in two cricket species via growth delays and physiological limits

**DOI:** 10.1101/2025.08.25.672160

**Authors:** Émile Vadboncoeur, Marie-Hélène Deschamps, Susan M. Bertram, Heath A. MacMillan

## Abstract

Understanding how chronic environmental stressors shape animal development is essential for predicting ecological responses and optimizing rearing systems. This perspective complements the use of short-term tolerance assays, which overlook the cumulative effects of sustained stress. Temperature and nutrition affect key life-history traits such as growth, development rate, and survival, which are closely tied to reproductive success and fitness. While both factors have been widely studied, their relative impacts aren’t clearly defined. We investigated how constant temperature (26– 41°C) and dietary protein-to-carbohydrate (P:C) ratio (0.15–2.18) influence development in two cricket species, *Acheta domesticus* and *Gryllodes sigillatus*. Growth trajectories were modelled using a unified-logistic equation to estimate asymptotic mass and relative growth rate. This approach captures the growth trajectory in a simplified and interpretable way, enabling comparisons across treatments. Asymptotic mass was combined with developmental rate and survival to calculate a composite metric of developmental performance. Developmental performance peaked at 35°C but fell at thermal extremes due to delayed development (in cold) or reduced mass and survival (in heat). Diet had more modest effects. Performance was stable across most P:C ratios, declining only at extreme imbalances. Notably, the performance cost of the most unbalanced diets was comparable to a 4-5°C shift from thermal optimum. Our results demonstrate that temperature, more than diet, drives variation in developmental performance during *ad libitum* feeding. This integrative framework provides a robust approach to quantify environmental sensitivity, define performance limits, and guide us toward the mechanisms underlying those limits and/or performance trade-offs.

**Summary statement:** Temperature more strongly influences growth trajectories and developmental performance than diet, due to delayed development and reduced survival at thermal extremes.

## Introduction

Understanding how chronic environmental stressors shape insect development is critical for predicting ecological responses and optimizing mass rearing systems. Reproductive success depends on development, growth, and survival traits, and both temperature and diet (two of the most widely studied factors) can influence all three traits (Clissold and Simpson, 2015; Hardison et al., 2021; Johnson et al., 1992; Raynal et al., 2022; Sissener et al., 2021). Collectively, these traits govern the likelihood of reaching reproductive maturity, a fundamental determinant of fitness with broad applications in conservation, pest control and mass-rearing. The relative influence of temperature and diet on each of these traits, however, remains unclear.

While dietary manipulations are frequently tested over the full course of development, short-term exposures to temperature are often used to estimate optimal temperatures and upper and lower thermal limits, particularly through metrics such as critical thermal limits. Although convenient, these acute assays have been increasingly criticized for their limited ecological relevance (Desforges et al., 2023; Kingsolver and Umbanhowar, 2018; Leong et al., 2022; Ørsted et al., 2022). In contrast to diets, which are rarely thought of as acute stress, high temperatures may appear tolerable in short-term trials despite ultimately impairing survival, growth, or development (Abarca et al., 2024). Such survivor bias can obscure biologically meaningful thresholds and hinder comparisons across species or treatments with differing physiological sensitivities. Chronic exposure experiments, which assess performance over extended periods and across multiple traits, offer a more integrative understanding of thermal stress (Abarca et al., 2024; Kong et al., 2025b). By quantifying survival, development time, and mass under constant temperature conditions, we can better identify sublethal thermal limits and compare their effects to those of dietary imbalance, contextualizing performance trade-offs across environmental gradients through full development.

Insect performance traits typically vary across environmental gradients, with low performance at extreme values and peak performance at intermediate values. Temperature effects on performance have long been a research focus, and thermal performance curves have been documented for numerous traits in many insect species (Alruiz et al., 2023; Magara et al., 2024; Rebaudo and Rabhi, 2018). For example, survival probability remains high across moderate broad temperature but declines steeply near species-specific critical thermal limits (**Figure 1A**; Abarca et al., 2024; Régnière et al., 2012). Developmental rates follow a typical thermal performance pattern, increasing with temperature up to an optimum and then dropping sharply (**Figure 1A**; Gibert and De Jong, 2001; Lamb and Gerber, 1985; Morales-Ramos et al., 2024). Critical limits are often defined as the temperature at which physiological rates, such as development, growth, or survival, reach zero (Angilletta, 2009). Mass at adulthood is not a trait that thermal performance curves can accurately describe (Angilletta, 2009), but is important to reproductive fitness (Kingsolver and Pfennig, 2004; Peters, 1986). For many ectotherms, body size at adulthood follows the temperature-size rule, where animals grow larger at lower temperatures (Atkinson, 1994). However, some taxa grow largest at intermediate or higher temperatures, including orthopterans, like crickets (**Figure 1A**; Kong et al., 2025b; Whitman, 2008).

**Figure 1.**
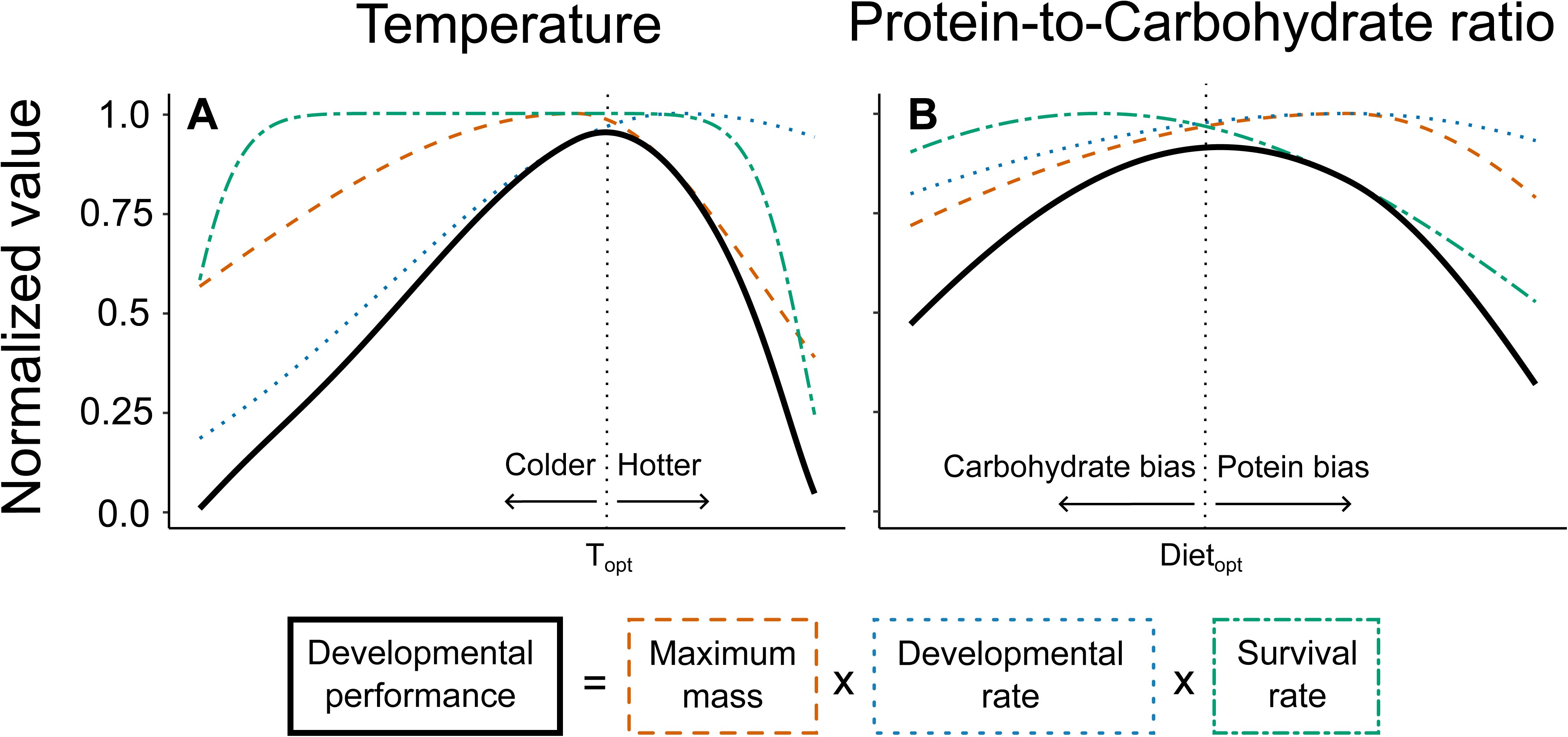
Conceptual framework illustrating how temperature (A) and protein-to-carbohydrate (P:C) ratio (B) influence key life-history traits and how these traits combine into a composite metric of developmental performance. Each panel displays two normalized life-history traits: maximum mass (orange dashed line) and developmental rate (blue dotted line), and survival rate (green dot-dashed line), which multiply together to form a developmental performance metric (solid black line). The existing literature guides the shape of these predictive curves for insects with a specific focus on Orthoptera when appropriate (see **Table S1** in the supplementary material). In panel A, survival remains high over a broad thermal range before declining sharply at upper and lower limits; developmental rate increases with temperature up to an optimum, then stabilizes or declines; and maximum mass tends to peak at intermediate to high temperatures in orthopterans, decreasing at both extremes, with a steeper decline at high temperatures. Developmental performance peaks at an optimal temperature (T_opt_), where the combined product of traits is maximized. At higher temperatures, performance declines steeply due to reduced survival and mass, while at cooler temperatures, it declines more gradually, primarily due to slower development despite high survival and mass. Thermal limits in insects are generally well defined by temperatures at which animals cannot survive over short to medium time frames. In panel B, survival peaks on carbohydrate-rich diets and declines with increasing protein content; developmental rate increases with protein content, peaking at balanced or protein-biased P:C ratios; and maximum mass follows a similar trend, reaching its highest values near balanced or protein-biased diets and decreasing under both protein- and carbohydrate-dominated conditions. Developmental performance peaks at an optimal macronutrient ratio (Diet_opt_), where survival, growth, and development are simultaneously maximized. At carbohydrate-biased extremes, performance declines due to slower development and smaller body size, while at high protein levels, elevated mortality constrains performance. Unlike temperature, dietary critical limits are less abrupt, but extreme macronutrient imbalances can still restrict developmental success. By integrating multiple traits, this metric allows direct comparison of how different environmental parameters shape overall developmental outcomes and reveals which exerts greater influence across the tested range. We hypothesize that the amplitude of variation in developmental performance across the dietary gradient will be smaller than across the thermal gradient, reflecting the more stringent physiological constraints imposed by extreme temperatures.

In contrast to temperature, the effects of diet on performance are more difficult to generalize due to variation in ingredient composition. However, a standardized approach for studying diet effects on performance in insects involves manipulating the ratio of protein to carbohydrate (P:C) (Raubenheimer and Simpson, 1999; Simpson and Raubenheimer, 1995). As protein content increases, survival tends to decrease (**Figure 1B**; (Bouchebti et al., 2022; Dussutour & Simpson, 2012; Lee, 2015; Lee et al., 2008a; Muzzatti et al., 2024; Nicholls et al., 2021), while developmental time is typically shortest on balanced or slightly protein-biased diets (**Figure 1B**; Kim et al., 2020; Lee, 2015; Muzzatti et al., 2024; Roeder and Behmer, 2014). Adult mass is similarly maximized under balanced to slightly protein-biased conditions (**Figure 1B**; (Jang and Lee, 2018; Kaewtapee et al., 2024; Muzzatti et al., 2024; Roeder and Behmer, 2014). Performance declines at the extremes of the P:C spectrum (approaching 1:0 or 0:1; Muzzatti et al., 2024; Roeder and Behmer, 2014). But, unlike temperature, clear physiological critical limits for P:C ratios aren’t well-defined, in part due to the impressive ability of insects to regulate intake and compensate for imbalanced diets (Behmer, 2009).

Traditionally, traits such as development time, maximum mass, and survival rate have been examined independently to assess the effects of environmental conditions on organismal performance. However, evaluating these traits in isolation can obscure underlying trade-offs and interactions. For example, larger body sizes achieved through prolonged development may be disadvantageous in season-limited environments (Angilletta et al., 2004; Nufio et al., 2025). To address this, we adopted a formula proposed by Muzzatti et al. (2024) that integrates these three traits into a single measure of individualized yield: the amount of biomass produced per unit time, conditional on survival. While Muzzatti et al. (2024) refer to this metric as yield, reflecting its relevance to farming contexts, we refer to it here as developmental performance, as it captures multiple metrics of performance during the developmental period. In our revised version of the metric, we also normalize mass and developmental rate to the mean of the control group to facilitate comparison between species and sexes that reach different adult sizes. Plotting developmental performance across a range of parameter values (**Figure 1A** and **1B**) allows clearer identification of optima and limits and enables more comprehensive comparisons between parameters such as temperature and diet. This approach addresses calls for more holistic metrics of performance that reflect ecological fitness (Abarca et al., 2024).

Understanding how environmental conditions affect insect development is key not only for predicting population dynamics in ecological contexts (Camacho et al., 2024; Overgaard et al., 2014), but also for improving outcomes in the context of insect mass rearing for farming (Kong et al., 2025b; Li et al., 2023; Muzzatti et al., 2024), pest management (Johnson et al., 1992; Nyamukondiwa et al., 2013; Quispe-Tarqui et al., 2021), and mitigating disease spread (Lee et al., 2008b; Wojda, 2017). Thermal sensitivity of insect traits is frequently studied to anticipate range shifts, optimize rearing conditions, or inform control strategies (Kong et al., 2025b; Pinkert et al., 2025). While insect diets are also of interest for these purposes (Clissold and Simpson, 2015; Muzzatti et al., 2024), diet effects are rarely evaluated at several temperatures. Identifying which environmental axes have the greatest influence on developmental performance could guide more targeted decision-making.

In this study, we examine two cricket species, *Gryllodes sigillatus* and *Acheta domesticus*, to assess how a range of temperatures and dietary protein-to-carbohydrate (P:C) ratios affect developmental time, growth, and survival, and a composite metric of developmental performance that integrates all three traits. Our design allows us to quantify the relative magnitude of environmental effects on each trait and also combine them in a single developmental metric. We found that temperature had a strong and consistent influence on developmental performance, primarily by accelerating or delaying development and reducing adult mass at thermal extremes. In contrast, diet only affected performance at the most unbalanced P:C ratios: very low protein diets reduced adult mass, while very high protein diets reduced survival. Notably, the largest diet-induced decline in performance was equivalent to only a 4-5°C deviation from the thermal optimum. Together, these findings demonstrate that temperature is a dominant environmental driver of developmental performance in crickets, acting mainly through its effects on developmental timing.

## Materials and methods

### Experimental diets

Oligidic diets were formulated using practical ingredients from a nearby feed mill (Campbellford Mill, Campbellford, ON) to better reflect natural or applied feeding conditions. Proportions of ingredients in the experimental diets were chosen to achieve different protein-to-carbohydrate ratios (**Table 1**). We opted for this approach over the geometric framework (Raubenheimer and Simpson, 1999; Simpson and Raubenheimer, 1995) as this method lacks a straightforward means of comparison with temperature, a continuous variable that cannot be diluted in the same way as diet. To standardize particle size and prevent selective feeding, all ingredients were sifted to remove particles larger than 1 mm, and any remaining larger particles were ground using a mortar and pestle until they passed through a 1 mm mesh.

**Table 1.** Formulation and proximal composition of oligidic diets having different protein-to-carbohydrate ratios.

| P:C | 0.15 | 0.22 | 0.45 | 0.83 | 1.36 | 1.75 | 2.18 |
| --- | --- | --- | --- | --- | --- | --- | --- |
| Ingredients (%) |  |  |  |  |  |  |  |
| Corn | 86 | 73.8 | 47 | 28.8 | 10 | 3 | 2 |
| Soymeal | 5 | 10 | 33 | 29 | 22 | 13 | 3 |
| Canola meal | 2 | 5 | 5 | 5 | 5 | 5 | 5 |
| Corn gluten | 4.3 | 8 | 10.8 | 25 | 50.8 | 62.8 | 62.6 |
| Fishmeal | 0.5 | 1 | 2 | 10 | 10 | 14 | 25 |
| Hog pro mix<sup>1</sup> | 2.2 | 2.2 | 2.2 | 2.2 | 2.2 | 2.2 | 2.2 |
| Composition (% , dry basis) |  |  |  |  |  |  |  |
| Crude proteins | 12.5 | 16.8 | 26.9 | 39.6 | 51.4 | 56.8 | 59.7 |
| Carbohydrates | 80.3 | 75.0 | 62.0 | 49.0 | 37.6 | 32.4 | 27.4 |
| Crude lipids | 1.8 | 1.8 | 2.3 | 2.8 | 2.7 | 2.7 | 3.8 |
| Crude fibers | 2.4 | 3.0 | 3.4 | 2.8 | 2.5 | 2.0 | 1.8 |
| Ash | 3.0 | 3.4 | 5.3 | 5.8 | 6.0 | 6.2 | 7.3 |
| Energy (kcal/g) | 3.9 | 3.9 | 4.0 | 4.3 | 4.5 | 4.7 | 4.7 |
| Dry Matter | 88.5 | 88.7 | 89.1 | 90.4 | 90.7 | 91.0 | 91.4 |

<sup>1</sup> Mix of minerals and vitamins for swine feed produced by Bio-Ag Consultants and distributors (Wellesley, On, Canada)

Proximal analyses were conducted for all diets (**Table 1**). Ash content was measured by incinerating diet samples at 600°C for 13 h according to AOAC, 942.05 (Thiex et al., 2012) using a Lindberg/Blue M 1100 C Box Furnace (ThermoScientific, Burlington, ON). Proteins were determined using the Dumas method (high-temperature combustion) in accordance with AOAC 992.23, using a FP828p analyzer (LECO Corporation, Mississauga, ON). Total lipids were extracted with diethyl ether using an XT15 extractor (ANKOM Technology, Macedon, NY, USA) according to AOCS Am 5-04. Crude fiber content was measured by sequential digestion with 0.255N H_2_SO_4_ and 0.313N NaOH using ANKOM filter bags, following AOCS Ba-6a-05, and analyzed with an ANKOM 200 fiber analyzer (ANKOM Technology, Macedon, NY, USA). Total carbohydrate content was estimated by subtracting the sum of measured ash, protein, lipid, and fiber values from 100%. Energy content was determined via bomb calorimetry using a 6400 Automatic Isoperibol Calorimeter (Parr Instrument Co., Moline, IL, USA), following the manufacturer’s instructions.

### Cricket rearing

Our *G. sigillatus* colony originated from eggs from Entomo Farms (Norwood, ON, Canada) and is maintained at Carleton University (Ottawa, ON, Canada), where 20 families are kept for laboratory use. One week after reaching adulthood, the crickets lay eggs in peat moss over two days and are then incubated until hatching. Eggs of *A. domesticus* came from Aspire Food Group (London, Ontario, Canada). Adults laid eggs in peat moss overnight, after which the moss containing the eggs was transferred into an insulated container with heat packs and shipped to Carleton University. Eggs arrived within 24 hours of being laid, then returned to an incubator to complete development. All eggs were incubated at 32°C under a 14 h light:10h dark photoperiod in moistened peat moss. The experiment started one day after hatching (11 and 9 days for *G. sigillatus* and *A. domesticus*, respectively). One-day post-hatching crickets were placed in individual condiment container (96 mL, 7 cm diameter × 3 cm depth, **Figure S1**) that contained a piece of egg carton for shelter. Water was provided via a 1.5 mL microcentrifuge tube sealed with dental cotton, allowing crickets to drink. Feed was provided *ad libitum* in a small plastic ink cap (1.7 cm diameter × 1.4 cm depth) glued to a 50 mm petri dish. The feed dishes and water tubes were swapped every two and three days, respectively. The crickets were kept on a 14 h light:10 h dark photoperiod at a relative humidity ranging from 65 to 85% for all incubators. Temperature and humidity were recorded every 30 min using an environmental logger (Inkbird, Shenzhen, China).

### Experimental treatments

The experimental design was established to facilitate the comparison between the effects of temperature and diet by linking treatments using a control (32°C 0.83 P:C) for both experiments and species. Each treatment combination included 30 individual crickets per species (unless mentioned otherwise below), resulting in a total sample size of 700 individuals.

Crickets of both species were reared on a control diet (balanced diet of 0.83 P:C) at different temperatures (26, 29, 32, 35, 38, 41°C). The control diet was selected because it approximates the intake target observed in self-selection trials for *G. sigillatus* (1.05 P:C; Muzzatti et al., 2024). While no specific intake target has been documented for *A. domesticus*, orthopterans generally select balanced diets (Behmer, 2009). All temperature treatments were conducted using a customized array of mini-incubators (Vevor Inc., Shanghai, China).

Crickets of both species were reared on different diets at a constant temperature of 32°C. For *G. sigillatus*, the temperature was chosen because it lies at the lower end of the thermal range that maximizes most life-history traits (Kong et al., 2025b). For *A. domesticus*, 32°C is slightly higher than the 30°C proposed by Clifford and Woodring (1990), but still within the optimal range (28 - 35°C; Clifford et al., 1977).

Seven (0.15, 0.22, 0.45, 0.83, 1.36, 1.75 and 2.18; P:C) and five (0.15, 0.22, 0.45, 0.83 and 1.75; P:C) experimental diets were tested in *G. sigillatus* and *A. domesticus,* respectively. For the *G. sigillatus*, diets from 0.45 to 2.18 were divided evenly across four mini-incubators, with six individuals per treatment per incubator. The two most carbohydrate-biased diets (0.15 and 0.22) were tested in a follow-up trial, housed in three additional incubators. Each of these incubators contained ten individuals per new diet, plus five individuals from the 0.45 and 0.83 groups to ensure consistency with the earlier trial. For *A. domesticus*, treatments were conducted in a large incubator (Thermo Fisher Scientific Inc., Massachusetts, United States).

### Data collection

Three times a week, the survival status was noted, and the position of each container was randomized to avoid positional effects. Every seven days, surviving individuals were weighed using an analytical scale (Mettler Toledo AB135-S, ON, Canada). Sex (when discernible at antepenultimate instar) and instar (based on moult counts) were recorded according to Kong et al., (2025a). The experiments were stopped at six weeks (42 days) as it encompassed the developmental period of the two species.

### Modelling individual growth and development

Data from insects are increasingly modelled to observe the effects of diets (Kaewtapee et al., 2024; Sripontan et al., 2020) or temperature (Grunert et al., 2015) on growth. Here, we fitted models to the growth and development trajectories of each individual cricket and compared the resulting parameters within species across treatments to better understand how diet and temperature affect these traits. To evaluate how mass changed over time, we modelled individual growth curves using three-parameter sigmoid unified-logistic regressions (Tjørve and Tjørve, 2017). Unified-logistic models were selected for their superior fit and interpretability across treatments when compared (R² values and visual inspection of residuals) to other commonly used sigmoid models describing insect growth (Grunert et al., 2015; Kaewtapee et al., 2024; Sripontan et al., 2020): U-Gompertz (n=555, 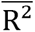 = 0.995), U-Bertalanffy (n = 476, 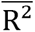 = 0.963) and U-Richards (n = 505, 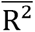 = 0.907). Since the Gompertz and Bertalanffy models tended to overestimate asymptotes, the unified-logistic regression (n = 579, 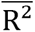 = 0.994) was chosen for its consistent convergence and superior fit, especially in estimating asymptotic mass.

Unified-logistic model has two forms: one uses the inflection point, while the other uses the y-intercept to anchor the curve. Since our study measured growth from hatching, we used the intercept-based form to anchor the curve to the starting mass (i.e. hatching mass) of each cricket. We therefore used the unified-logistic formula as follows:

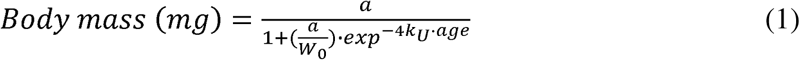

where a is the asymptotic mass (i.e., maximum body mass), W is the body mass at hatch (y-intercept), and k_U_ is the relative growth rate, which controls how quickly the curve approaches the asymptote (i.e., k_U_ = 0 is flat, k_U_ = ∞ is vertical). **Figure 2** illustrates how each parameter influences the growth curve.

**Figure 2.**
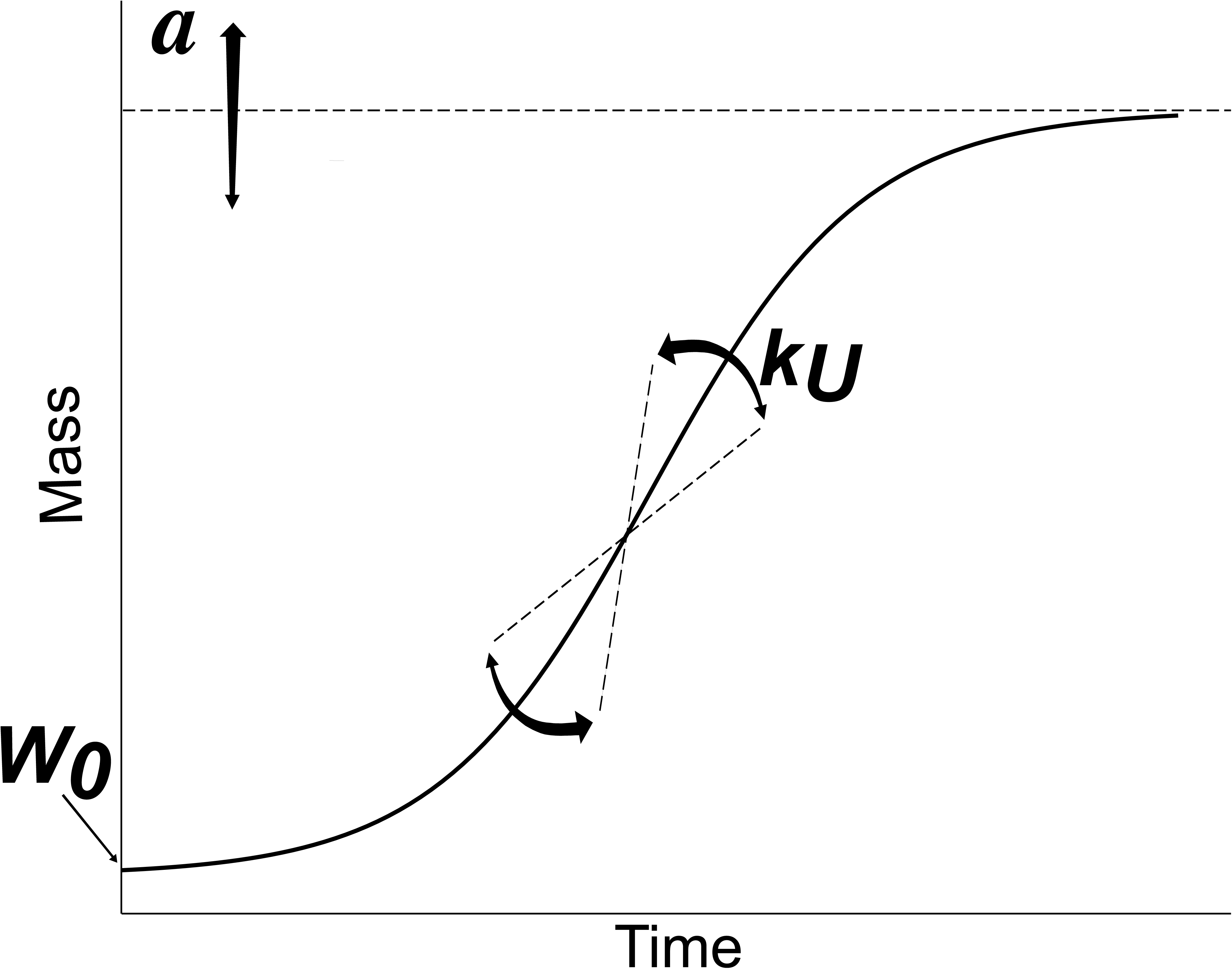
Example growth curve of a cricket, illustrating the three parameters of the unified logistic model (Equation 1). The asymptote (**a**) represents the estimated maximum body mass at the end of development. The intercept (**W_0_**) reflects the body mass at hatching. The parameter **k_U_** indicates the relative maximum growth rate, describing how quickly the curve approaches the asymptote. Together, these parameters characterize the full trajectory of growth from hatch to maturity.

Developmental time is typically measured as the number of days until reaching the final molt or a specified pre-adult instar (Magara et al., 2024; Morales-Ramos et al., 2024; Muzzatti et al., 2024). In this study, instar stage was recorded weekly, which did not allow us to precisely determine the day of adult emergence. Instead, we estimated developmental rate by fitting a linear regression to weekly instar data for each individual:

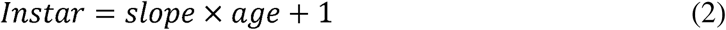

where the slope represents the rate of instar progression, and the intercept was fixed at 1, corresponding to the first instar at hatch. Only individuals surviving to at least week 5 were included in this analysis. An exponential model (n = 478, 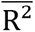 = 0.985) was also tested, but the linear model (n = 575, 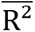 = 0.962) was preferred for its simplicity and convergence across individuals. Model fits were strong overall, with R^²^ values > 0.95 for growth and > 0.85 for development. Notably, this technique preserves variance in individual growth and development trajectories but also compounds errors from individual regressions when comparing mean parameters for the different experimental groups. With consistently strong model fits (high R^2^ values), we are confident that this error is minimized and that the parameters accurately represent the growth and development of our experimental animals. Examples of unified-logistic growth curves are shown in **Figure 3 and Figure S2**.

**Figure 3.**
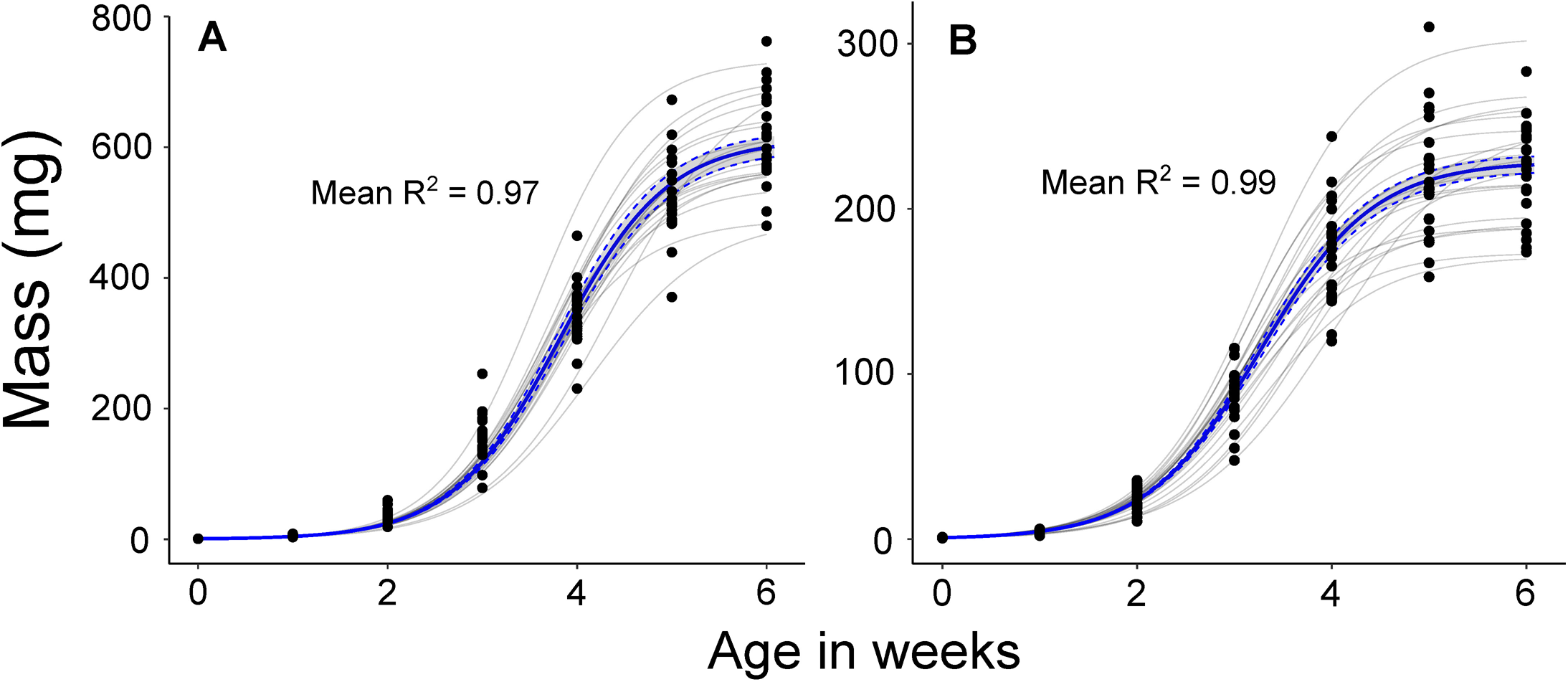
Examples of individual and average growth trajectories of female *A. domesticus* (A) and male *G. sigillatus* (B) reared at 32°C on a 0.83 protein-to-carbohydrate (P:C) diet. Black points represent weekly body mass measurements for each cricket. Grey lines show unified logistic model fits for each cricket, while the bold blue line represents the mean fitted curve with shaded 95% confidence interval. Mean R^2^ values indicate model fit across individuals within each group.

### Developmental performance

Developmental performance was calculated using the yield metric described by Muzzatti et al. (2024), which integrates survival, developmental rate, and adult body mass into a single value for every individual. The metric was adapted as follows:

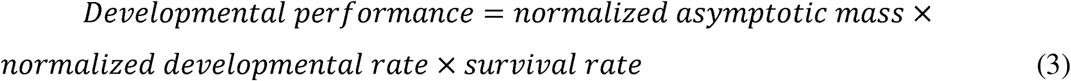

The asymptotic mass was derived from the unified logistic growth model (parameter a, see Section 2.4). The developmental rate is the slope of the linear regression through the increase of instar through time (see Section 2.4). Both asymptotic mass and developmental rate are normalized to control conditions (32°C; 0.83 P:C). Survival was calculated per treatment group and applied uniformly to all individuals within that group. It was defined as the proportion of individuals alive on day 42 relative to initial number of individuals in that group. This value, specific to each species and treatment group, was not normalized, as it already falls between 0 and 1 and reflects a biologically meaningful component of development.

### Modelling the different parameters

To model the effects of temperature and diet on developmental rate and performance, we fit thermal performance curves using models from the “rTPC” package in R. For temperature-dependent traits, several models were tested, and the best-fitting models were selected based on Akaike Information Criterion (AIC). For diet-dependent traits, only a quadratic model was tested, as used in nutritional geometry (Jensen et al., 2015; Lee et al., 2008a). For each trait, the model with the lowest AIC was visually inspected to ensure it captured the general shape of the response. The best-fitting model was then used to generate a continuous curve describing how each trait varied across the temperature or diet.

### Statistical analysis

All analyses used R 4.3.3 (R Core Team, 2024; https://www.r-project.org/). All figure data are reported as means ± sem. Given the goal of the current study to compare the thermal and dietary effects, species were not compared statistically; our analysis compared the different treatments and sexes. To fit nonlinear models, we used the “*nls_multstart*” function from the “*nls.multstart*” package to extract parameter estimates. All parameters were compared to the control group of crickets held at 32°C and 0.83 P:C. Survival data were analyzed using a Cox proportional hazards model with the “*coxph”* function in the *survival* package with species and treatment as predictors. Extracted growth and development parameters, as well as developmental performance, were analyzed using generalized linear models with a “*Gamma”* distribution, incorporating treatment, sex, and their interaction as fixed effects. The “*stepAIC”* function from the *MASS* package to identify the most parsimonious models. The selected models and their outputs can be found in **Table S2 to S11**. Every generalized linear model was examined to ensure that the residuals were normally distributed and independent.

## Results

### Developmental Rate: Temperature-Driven, Diet Stable

Developmental rate, measured as the slope of instar progression over time, was strongly influenced by temperature. Both *G. sigillatus* and *A. domesticus* developed faster at 35°C, with rates reaching approximately 1.8 instars per week, but declined at both cooler and warmer temperatures (**Figure 4A & C**). In stark contrast to temperature, diet had minimal effects on developmental rate. For *G. sigillatus*, no significant differences were observed across widely different diets **(Figure 4D**). For *A. domesticus*, only the most protein-biased (t = 3.14, p < 0.001) and most carbohydrate-biased (t = 2.43, p = 0.016) diets resulted in significantly slower development compared to the control (**Figure 4B).**

**Figure 4.**
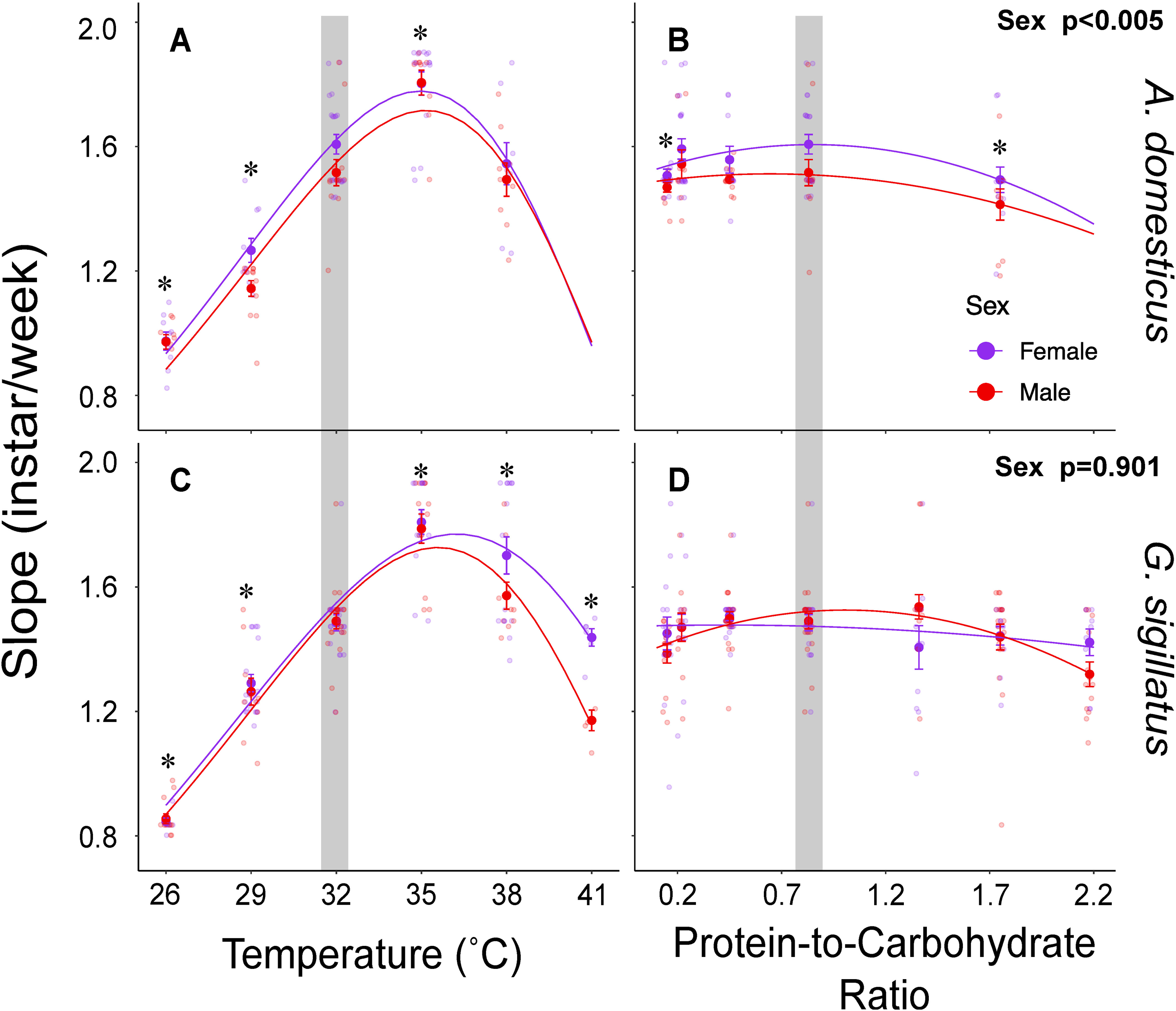
Developmental rate is strongly temperature-dependent but largely unaffected by diet. Developmental rate, measured as the slope (instars/week) of a linear regression fitted to weekly instar data for male (red) and female (purple) *A. domesticus* (left panels) and *G. sigillatus* (right panels). Panels A and C show developmental rates across temperatures (26, 29, 32, 35, 38, and 41°C), while panels B and D show rates across diets with varying in P:C (0.15 to 2.18). Each point represents an individual; lines show model predictions. The shaded region indicates the control condition (32°C and 0.83 P:C diet). Asterisks (*) denote significant differences (p < 0.05) from the control.

### Growth trajectories

#### Overall effects of temperature and diet on growth

Temperature and diet shaped overall growth trajectories of both species by influencing asymptotic mass and relative maximum growth rate (k_U_) (**Figure 5**). Temperatures above or below the thermal optimum reduced k_U_, flattening the growth curve and delaying its progression toward the asymptote, which itself also declined. For example, female *A. domesticus* reared at 32°C reached 99% of their asymptotic mass (612□±□15 mg) in 6.4□±□0.1 weeks, whereas those reared at 26°C reached 99% of a lower asymptote (432□±□41 mg) in 10.0□±□0.2 weeks. At 38°C, females reached 99% of their final mass more quickly (5.4□±□0.3 weeks), but the asymptote was less than half the control value (257□±□26 mg). Similar temperature-related trade-offs were observed across both sexes and species.

**Figure 5.**
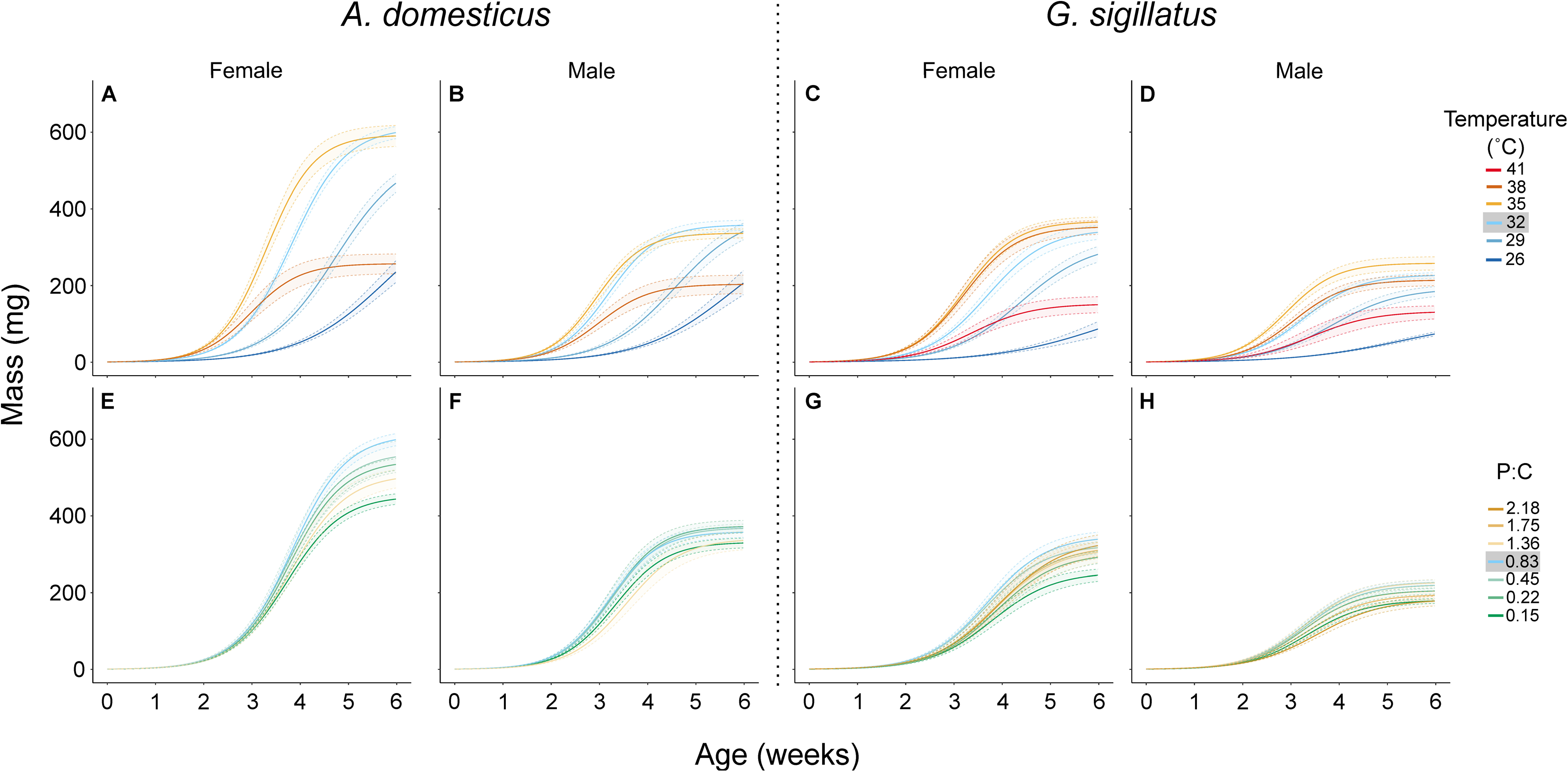
Modeled growth trajectories of female and male *A. domesticus* (left panels) and *G. sigillatus* (right panels) reared under different temperature (A–D) and dietary (E–H) treatments. Panels A–D show mass (mg) over time across six temperatures (26, 29, 32, 35, 38, and 41°C). Panels E–H show the effects of seven protein-to-carbohydrate (P:C) ratios (0.15 to 2.18) on growth at a constant temperature of 32°C. Solid lines represent logistic regression fits; shaded ribbons indicate standard error. Female and male data are shown in the left and right columns of each species panel, respectively.

In contrast, the effects of diet on growth trajectories were comparatively modest. For instance, female *A. domesticus* fed the lowest P:C diet exhibited a reduced asymptotic mass (453 ± 13 mg) but reached 99% of it in the same amount of time as those on the control diet (6.4 ± 0.1 weeks). Likewise, female *G. sigillatus* fed the lowest P:C diet reached an asymptote 27% lower than the control, but without a notable delay in timing. Other diets caused less than a 20% reduction in asymptotic mass and did not significantly affect timing. In summary, extreme diets primarily reduced final mass with minimal impact on growth duration, while suboptimal temperatures altered both the size attained and the speed at which it was achieved.

#### A. domesticus

Newly hatched *A. domesticus* nymphs had an average mass of 0.74□±□0.09 mg (n = 310), consistent with the initial mass (W_0_) estimated by the unified logistic model (0.74□±□0.09 mg). By the final measurement day, control females reached 612□±□15 mg, significantly larger than control males (359□±□14 mg; t = 13.79, p < 0.001). The model-predicted asymptotic masses closely matched these values, validating the logistic model, with estimated asymptotes of 612□±□15 mg for females and 359□±□14 mg for males. The relative maximum growth rate (k_U_) was 0.44□±□0.01 for females and 0.49□±□0.01 for males. These sex differences were statistically significant for both asymptotic mass (t = 13.79, p < 0.001) and k_U_ (t = –3.81, p < 0.001). All comparisons reported below are relative to the control group of the same sex reared at 32°C on the 0.83 P:C diet (**Figure 6**).

**Figure 6.**
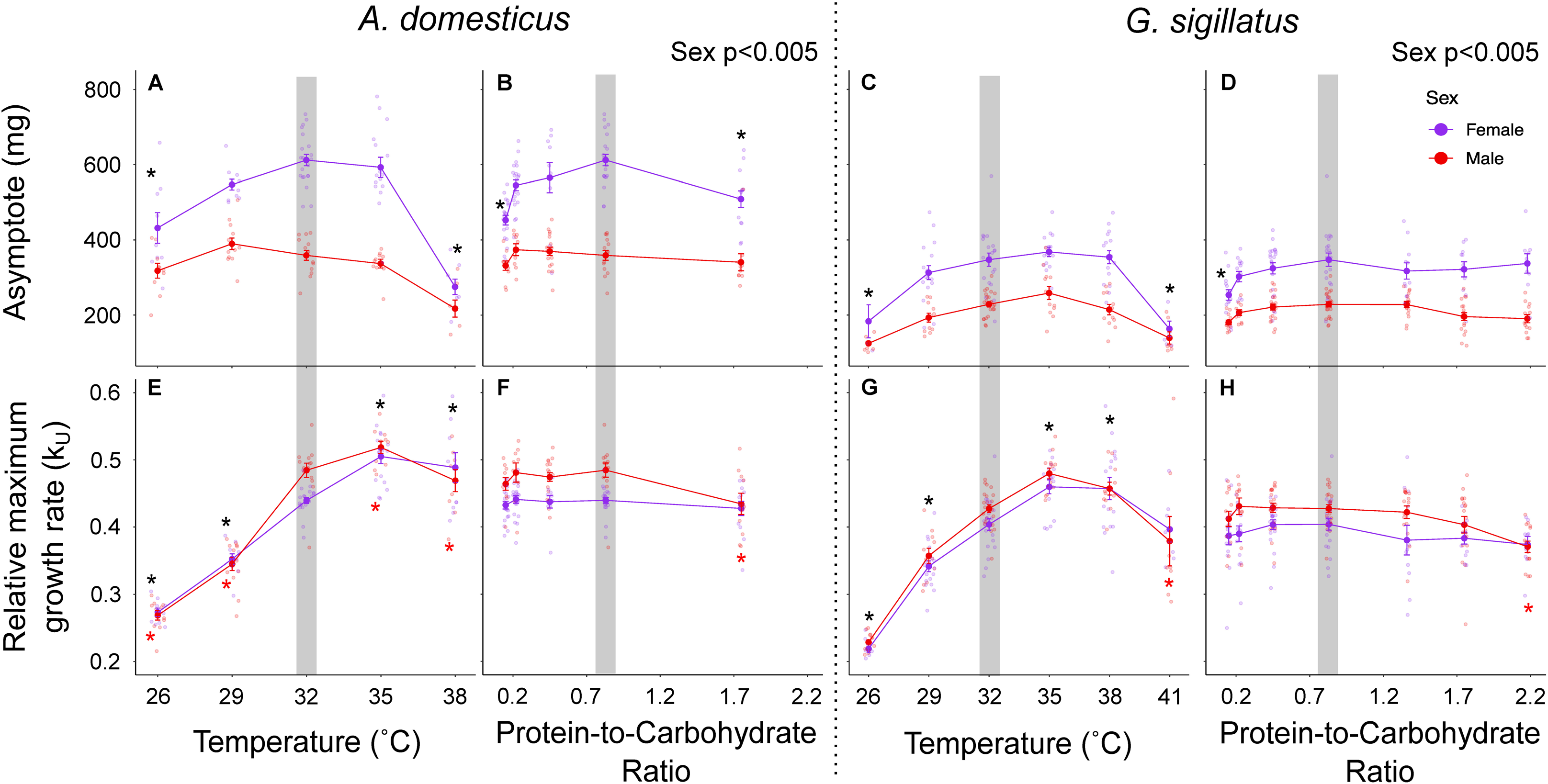
Unified logistic model parameters describing growth trajectories in male (red) and female (purple) *A. domesticus* (left panels) and *G. sigillatus* (right panels). Panels A–D show asymptotic mass (mg), and panels E–H show the relative maximum growth rate (k_U_), with effects shown across temperature (A, C, E, G) and dietary protein-to-carbohydrate (P:C) ratio (B, D, F, H). Each point represents an individual cricket reared from one day post-hatch to day 42 under a single treatment. The shaded region represents the control condition (32°C and 0.83 P:C diet). Asterisks (*) indicate a significant difference (p < 0.05) from the control (black) and interactions between sex and temperature (red). Asymptote refers to the modelled adult body mass, and k_U_ represents the rate at which mass increases toward the asymptote.

High and low temperatures, as well as the most protein-biased diet, significantly reduced asymptotic mass. The relative maximum growth rate (k_U_) was affected by temperature: higher temperatures increased k_U_, while lower temperatures reduced it. Sex differences in growth rate were temperature-dependent; there was a significant interaction between sex and temperature for k_U_ at 26°C (t = 2.82, p < 0.005), 29°C (t = 2.54, p = 0.01), 35°C (t = 2.03, p < 0.04), and 38°C (t = 3.35, p < 0.005), which was driven by a lack of sex differences in k_U_ at those temperatures, unlike the control group at 32°C. There was also a significant interactive effect of sex and diet on k_U_, which was similarly driven by a loss of sex difference in this parameter on the highest-protein diet (t = -2.04, p = 0.04), although no other differences in k_U_ were found among diets. The interactive effects between sex and the highest temperature and protein content in feed for k_U_ appear to be driven by reduced growth rates of males.

#### G. sigillatus

Newly hatched *G. sigillatus* had an average mass of 0.85□±□0.17 mg (n = 390), closely matching the starting weight (W_0_) estimated by the U-logistic model (0.86□±□0.16 mg). At the end of the study, control females reached 331□±□18 mg, and males 223□±□6 mg. These final masses aligned with the model’s estimated asymptotes of 347□±□18 mg for females and 229□±□7 mg for males. Despite larger hatchling mass, *G. sigillatus* adults were smaller than *A. domesticus*, indicating divergent growth strategies. The relative maximum growth rate (k_U_) was 0.40□±□0.01 for females and 0.43□±□0.01 for males. As with *A. domesticus*, these sex differences in both asymptotic mass (t = 13.98, p < 0.001) and k_U_ (t = –2.70, p = 0.007) were statistically significant. All reported treatment differences are relative to control crickets reared at 32°C and fed a 0.83 P:C diet (**Figure 6**).

Asymptotic mass of *G. sigillatus* declined to 144□±□24 mg at 26°C, 142□±□12 mg at 41°C, and 220□±□10 mg on the lowest-protein diet. k_U_ increased at 35 and 38°C but returned to baseline at 41°C and declined under cooler conditions. There was a significant interactive effect of sex and both temperature (t = 3.55, p < 0.001 at 41°C) and diet (t = 2.16, p = 0.031 for the lowest-protein diet) on k_U_, indicating that *G. sigillatus* males are more sensitive to nutritional and thermal stress than females.

### Survival

The survival rates in the control group (32°C and 0.83 P:C) were 97.8% for *G. sigillatus* and 87.5% for *A. domesticus* (**Figure 7**). Survival was only significantly reduced at temperature and dietary extremes, specifically under high temperatures and on the most protein- or carbohydrate-biased diets. For *A. domesticus*, survival at 41°C dropped to 0% by day 16. This rapid mortality suggests a critical upper thermal limit near 41°C for A. domesticus. By day 42, survival was 33% at 38°C and 77% on the 1.75 P:C diet. *G. sigillatus* exhibited greater thermal tolerance, maintaining 27% survival at 41°C on day 42, and survival on the 0.22 and 2.18 P:C diets was 83% and 73%, respectively.

**Figure 7.**
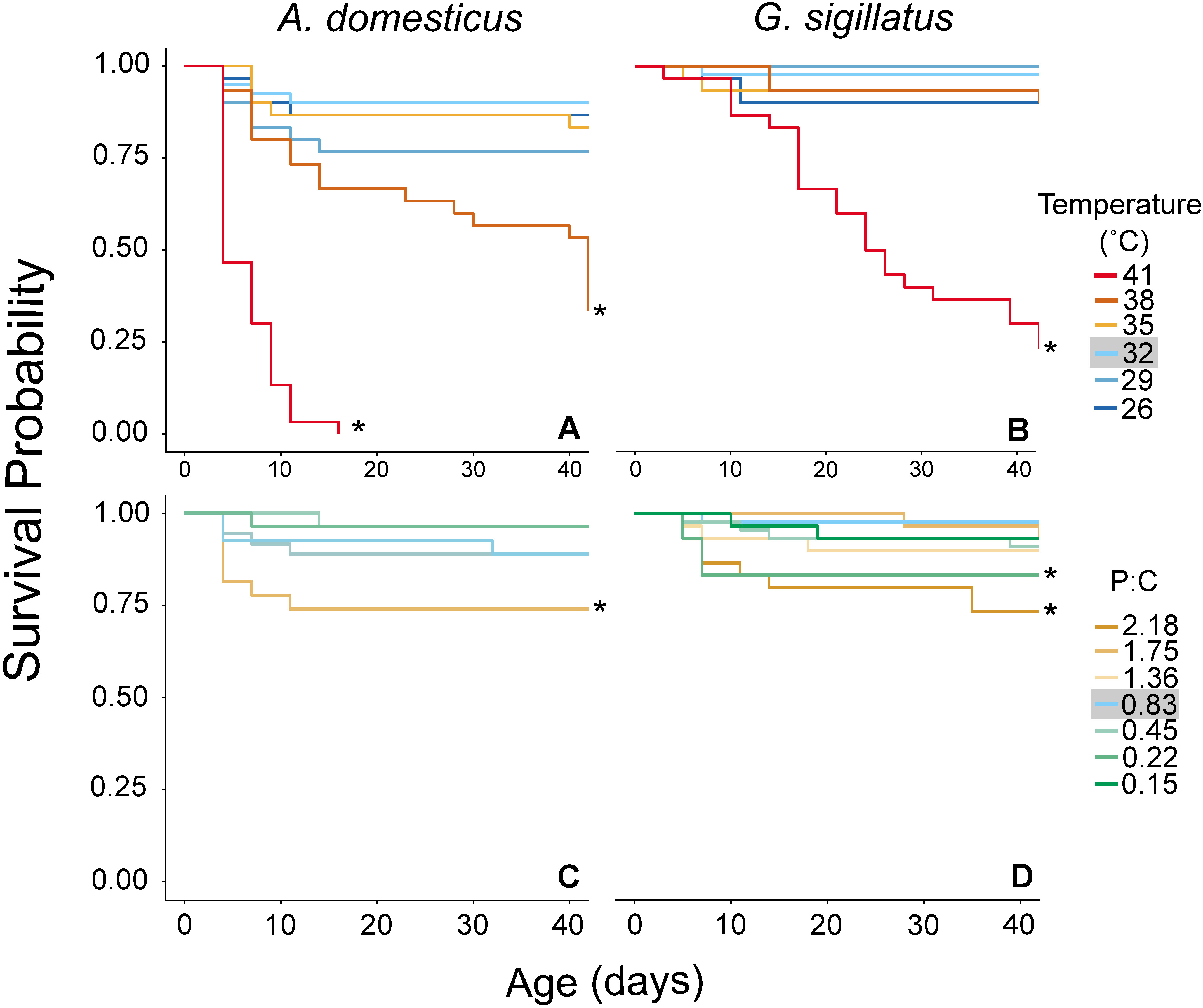
Temperature and diet both influence survival, but temperature effects are more severe. Survival probability of *A. domesticus* (A, C) and *G. sigillatus* (B, D) over 42 days under different temperature (A, B) and diet (C, D) treatments. Panels A and B show survival at six constant temperatures (26, 29, 32, 35, 38, and 41°C). Panels C and D show survival on seven experimental diets varying in protein-to-carbohydrate (P:C) ratio (0.15, 0.22, 0.45, 0.83, 1.36, 1.75, and 2.18). Asterisks (*) indicate a significant difference (p < 0.05) compared to the control group (32°C, 0.83 P:C diet; grey). Each group began with 30 individuals, except for the 0.83 and 0.45 P:C treatments at 32°C, which began with 45 individuals.

### Developmental Performance: Integrating Growth, Survival and Timing

Developmental performance was more strongly impacted by temperature than diet. Performance peaked at 35°C and declined at both thermal extremes, as a consequence of different traits (**Figure 8**), highlighting the asymmetric nature of thermal stress: low temperatures delayed development, while high temperatures reduced mass and survival. Dietary effects on developmental performance were more modest, with reductions observed only at the most extreme P:C ratios, specifically, the highest protein and carbohydrate levels. These findings reinforce the conclusion that temperature exerts a stronger influence on developmental performance than diet. Normalizing mass and growth rate to the mean of the control group rendered sex a non-significant influence on the developmental performance of both species, and we therefore excluded it from the analysis (*A. domesticus*: t=-0.059, p=0.95; *G. sigillatus:* t=0.787, p=0.43). For *A. domesticus*, developmental performance peaked at 35°C with the 0.83 P:C diet (0.91□±□0.03) and remained equivalent to control levels at 32°C on the same diet (0.88□±□0.02). From those temperature and dietary peaks, performance declined by 48% at 26°C (0.47□±□0.04); at 38°C, performance declined by 82% (0.17□±□0.01). Developmental performance only dropped by 19% (0.73□±□0.03) on the 0.15 P:C diet and by 30% (0.63□±□0.04) on the 1.75 P:C diet. Thus, the maximum change in developmental performance due to temperature exceeded that due to diet. For *G. sigillatus*, developmental performance peaked at 35°C with the 0.83 P:C diet (1.26□±□0.05). For the dietary range, control crickets at 32°C with the 0.83 P:C diet had the highest developmental performance (0.98□±□0.03) 22% lower than 35°C. Performance declined by 91% to 0.11□±□0.01 at the highest temperature and by 79% to 0.27□±□0.03 at the lowest temperatures; performance declined by 36% to 0.68□±□0.03 on the most carbohydrate-biased diet and by 40% to 0.59□±□0.04 on the most protein-biased diets. Temperature, therefore, impacted developmental performance more than diet. Indeed, across both species, the largest diet-driven decline in developmental performance was roughly equivalent to only a 4-5°C decrease from the thermal optimum.

**Figure 8.**
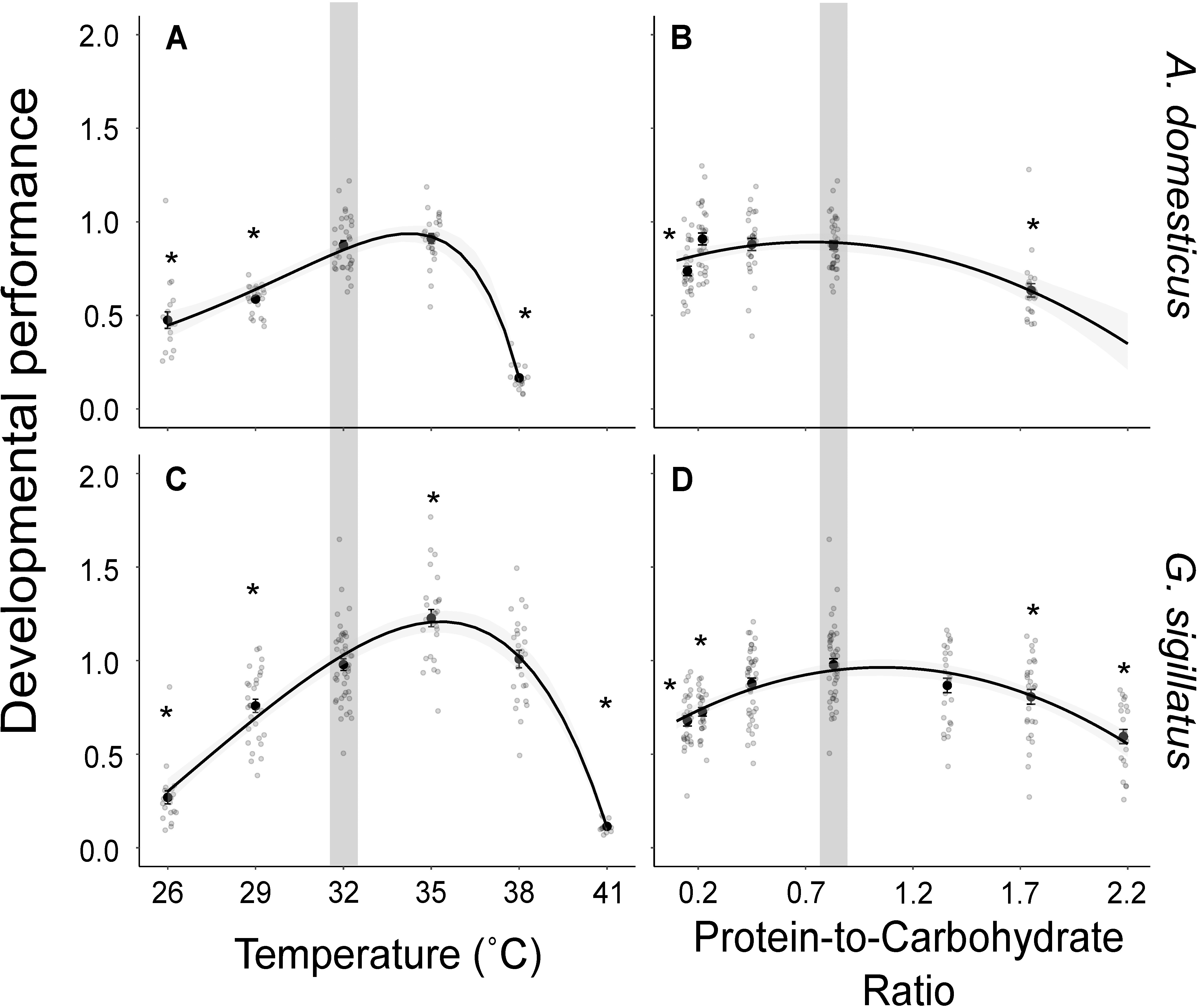
Developmental performance of *A. domesticus* (left panels) and *G. sigillatus* (right panels) reared from egg to adulthood under different temperature (A, C) and dietary protein content (B, D) conditions. Each point represents an individual cricket. Developmental performance was calculated by multiplying the ratio of asymptotic mass to the mean asymptotic mass of the control group by the survival probability of its treatment group and dividing by the number of weeks required to reach the 8th instar. The shaded region indicates the control condition (32°C and 0.83 P:C diet). Asterisks (*) denote significant differences (p < 0.05) from the control.

## Discussion

Understanding how animals respond to environmental stressors requires more than identifying physiological limits. It demands an integrated view of how stressors affect core developmental processes and the life-history trade-offs animals may need to make to complete development (Boggs, 2009; Zera and Harshman, 2001). While metrics like CT_max_ assays offer valuable insight into acute stressors, they often fail to capture the cumulative, sublethal effects of chronic exposure on growth, development and survival (Desforges et al., 2023; Kingsolver and Umbanhowar, 2018). Our study contributes to a broader effort in the animal performance literature to bridge this gap by quantifying how chronic exposure to two distinct stressors, temperature and dietary macronutrient balance, compares in shaping developmental performance. Using a composite performance metric, we show how chronic environmental conditions affect key developmental traits in two cricket species, advancing trait-based comparative frameworks essential for predicting organismal performance across contexts, from basic ecological theory to applied scenarios such as pest management and insect production systems (Kong et al., 2025b; Marshall et al., 2020; Muzzatti et al., 2024).

Here, we found that both temperature and diet influenced developmental outcomes and that temperature exerted a stronger and more consistent effect in two species of crickets. Developmental performance peaked at 35°C in both species and declined at both thermal extremes, but for different reasons: low temperatures delayed development while high temperatures reduced mass and survival. Diet, by contrast, only affected performance at the most unbalanced P:C ratios through effects on mass and survival. Notably, the largest performance reduction due to diet (from 0.83 to 0.15 P:C) was equivalent to only a 4-5°C deviation from the thermal optimum, indicating that temperature is the dominant environmental driver of developmental performance under *ad libitum* feeding.

The shape of the thermal performance curves for the different traits measured in both *A. domesticus* and *G. sigillatus* showed similar patterns. We found parallel steep declines in the developmental performance curves at high temperatures, that were driven by reduced growth and mortality. Similar to other studies, lower temperatures resulted in smaller adults but did not drive mortality (Abarca et al., 2024; Kong et al., 2025b; Lamb and Gerber, 1985; Magara et al., 2024). Our findings suggest temperatures below 26°C should be used to test lower chronic thermal stress on these two species. Our findings align with previous work showing that thermal performance curves for rate-based traits are typically skewed, with sharp declines at upper thermal limits (Rebaudo and Rabhi, 2018; Shi and Ge, 2010). In both species, developmental rate and asymptotic mass peaked at intermediate to high temperatures and declined at thermal extremes, consistent with this pattern. Although asymptotic mass is not a rate-based trait (Angilletta, 2009), the final size of an insect is determined by the rate of biomass accumulation and the duration of that growth period. As a result, the temperature dependence of asymptotic mass likely reflects the combined effects temperature on growth rate and development duration (Forster and Hirst, 2012). Unlike many ectotherms that follow the temperature-size rule (Atkinson, 1994), orthopterans such as crickets tend to reach maximum size at intermediate to warm temperatures (Kong et al., 2025b; Magara et al., 2024; Tu et al., 2012; Whitman, 2008), suggesting the thermal sensitivities of growth and development are closely aligned in this group (Forster and Hirst, 2012).

The range of P:C values we tested in this study had a comparatively limited effect on developing crickets. This was evident across all measured traits, including survival, growth, and development (**Figures 4–7**). Similar results have been reported in other herbivorous insects, where performance is maintained despite variation in dietary macronutrient balance (Deans et al., 2015; Deans et al., 2022; Lee et al., 2006; Muzzatti et al., 2024). Insects can mitigate suboptimal nutrient intake through a combination of behavioural and physiological strategies. While behavioural compensation, such as selecting alternative foods to balance intake, is common (Behmer, 2009), our crickets were restricted to a single diet. Crickets might have behaviourally compensated by increasing their food consumption, but this necessarily results in overconsumption of the non-limiting nutrient, which can come with metabolic or toxic costs (Behmer, 2009), discussed further below.

We argue that high temperatures induce high inefficiency realized over a short developmental period, while low temperatures induce modest inefficiency that extends over a prolonged period. These two scenarios culminate in similarly diminished growth outcomes. Both temperature extremes reduced asymptotic mass, but only low temperatures delayed the time required to reach adult mass. The highest temperatures tested (38°C for *A. domesticus* and 41°C for *G. sigillatus*) led to dramatic decreases in asymptotic mass but only modest reductions in relative maximum growth rate. In contrast, cooler temperatures reduced both relative maximum growth rate and final body size. Metabolic rate increases with temperature (Lachenicht et al., 2010; Roe et al., 1980), and so do growth rates, up to an optimal point beyond which performance declines (Bjørge et al., 2018). At these high temperatures, the efficiency with which energy is converted to mass is low, and animals tend to mature smaller (Bjørge et al., 2018), suggesting an inability to put resources towards growth because of increased maintenance costs associated with high temperature (González-Tokman et al., 2020). Lower temperatures similarly induce low energy conversion efficiency, associated with reduced growth rates (Bjørge et al., 2018), possibly indicating similar increased physiological costs, but likely for different reasons. Low temperatures reduce metabolic rate (Lachenicht et al., 2010; Roe et al., 1980), but also slow developmental processes (Booth and Kiddell, 2007; Magara et al., 2024), thereby extending the duration over which maintenance costs accumulate. Neither metabolic rate nor development time scales linearly with temperature, so the mismatch between energy availability and demand is greater further from the thermal optimum (Marshall et al., 2020). These patterns align with the broader framework of energetic allocation trade-offs (Mauritsson and Jonsson, 2023; Zera and Harshman, 2001), which propose that organisms balance limited energetic resources among competing demands, typically growth, maintenance, reproduction, and survival. At thermal extremes, resources are disproportionately reallocated toward maintenance and stress mitigation rather than somatic growth, leading to reduced asymptotic mass. Such trade-offs are central to life-history theory and have been observed in diverse ectotherms, including fish (Audzijonyte and Richards, 2018) and insects (Boggs, 2009) under environmental stress.

A similar accumulation of inefficiencies may explain the reduction in asymptotic mass under the most unbalanced diets. Crickets fed low-protein diets must consume more food to meet protein requirements (Simpson and Abisgold, 1985), potentially increasing the metabolic costs of feeding, digestion, and nutrient processing (Deans and Hutchison, 2023). Excess carbohydrate intake has also been linked to “wastage respiration”, an increase in CO production that does not contribute to growth (Zanotto et al., 1997). In addition, low P:C diets promote lipid accumulation in insects (Clark et al., 2014; Lee, 2015; Lee et al., 2015), which may advance the onset of moulting through size-sensing mechanisms (Nijhout and Callier, 2015). Therefore, the resulting individuals would be smaller if moulting is triggered before sufficient protein reserves are acquired to support maximum post-moult size.

High-protein diets may also reduce final mass, but through different costs. Surplus dietary protein is catabolized via gluconeogenesis (Zanotto et al., 1993), leading to nitrogenous waste excreted primarily as uric acid in insects (Weihrauch and O’Donnell, 2021). Because uric acid excretion is an active, energetically costly process, diets rich in protein may siphon off energy that could otherwise be allocated to growth, limiting maximum body size.

These patterns highlight how asymptotic mass and relative growth rate respond to different physiological constraints across thermal and dietary gradients. While the mechanisms driving reduced growth under extreme conditions likely differ, ranging from temperature-dependent metabolic inefficiencies to nutrient-specific processing costs, our modelling framework provides a powerful tool to disentangle their effects. By isolating growth rate and asymptote as independent parameters and reconstructing full growth trajectories, this approach allows researchers to infer the relative contributions of developmental timing, energy conversion efficiency, nutrient processing or any other possible mechanism to observed size outcomes. As such, growth modelling summarizes complex biological responses in a simplified and interpretable way and offers a pathway to develop and test mechanistic hypotheses about how environmental stressors reshape energy allocation strategies. This capacity makes it broadly applicable to both ecological research and applied efforts in insect rearing and performance optimization.

## Conclusion

Our results demonstrate that temperature exerts a stronger influence than diet on insect developmental performance. Performance peaked at 35°C in both species and declined at cooler and warmer temperatures, due to delayed development, and elevated mortality and reduced mass, respectively. Maximum mass was also reduced at temperature extremes. Diet composition had comparatively modest effects, with performance maintained across a broad range of protein-to-carbohydrate ratios and only declining under the most unbalanced conditions.

By parameterizing growth trajectories and integrating developmental rate, maximum mass, and survival into a unified performance metric, we propose a framework for assessing environmental sensitivity to different parameters. This approach may be particularly useful for comparing species with different life-history strategies or ecological niches and captures the developmental consequences of sustained environmental stress, providing a complementary lens to acute stress tolerance assays (Abarca et al., 2024; Kingsolver and Umbanhowar, 2018). Our results emphasize the value of multi-trait performance metrics in environmental research and offer a robust framework for future studies on environmental stress and life-history evolution.

## Acknowledgements

We want to thank Martha Paola Rivera Rodriguez for the proximal analysis of the diets at Université Laval. We also want to thank Aspire Food Group, which provided the eggs of *A. domesticus* used for this study.

## References

Abarca, M., Parker, A. L., Larsen, E. A., Umbanhowar, J., Earl, C., Guralnick, R., Kingsolver, J. and Ries, L. (2024). How development and survival combine to determine the thermal sensitivity of insects. PLOS ONE 19, e0291393.

Alruiz, J. M., Peralta-Maraver, I., Bozinovic, F., Santos, M. and Rezende, E. L. (2023). Temperature adaptation and its impact on the shape of performance curves in Drosophila populations. Proc. R. Soc. B Biol. Sci. 290, 20230507.

Angilletta, M. J. (2009). Thermal adaptation: a theoretical and empirical synthesis. Oxford: Oxford University Press.

Angilletta, M. J., Jr., Steury, T. D. and Sears, M. W. (2004). Temperature, Growth Rate, and Body Size in Ectotherms: Fitting Pieces of a Life-History Puzzle1. Integr. Comp. Biol. 44, 498–509.

Atkinson, D. (1994). Temperature and Organism Size—A Biological Law for Ectotherms? In Advances in Ecological Research, pp. 1–58. Elsevier.

Audzijonyte, A. and Richards, S. A. (2018). The Energetic Cost of Reproduction and Its Effect on Optimal Life-History Strategies. Am. Nat. 192, E150–E162.

Behmer, S. T. (2009). Insect Herbivore Nutrient Regulation. Annu. Rev. Entomol. 54, 165–187.

Bjørge, J. D., Overgaard, J., Malte, H., Gianotten, N. and Heckmann, L.-H. (2018). Role of temperature on growth and metabolic rate in the tenebrionid beetles Alphitobius diaperinus and Tenebrio molitor. J. Insect Physiol. 107, 89–96.

Boggs, C. L. (2009). Understanding insect life histories and senescence through a resource allocation lens. Funct. Ecol. 23, 27–37.

Booth, D. T. and Kiddell, K. (2007). Temperature and the energetics of development in the house cricket (Acheta domesticus). J. Insect Physiol. 53, 950–953.

Bouchebti, S., Cortés-Fossati, F., Vales Estepa, Á., Plaza Lozano, M., S. Calovi, D. and Arganda, S. (2022). Sex-Specific Effect of the Dietary Protein to Carbohydrate Ratio on Personality in the Dubia Cockroach. Insects 13, 133.

Camacho, A., Rodrigues, M. T., Jayyusi, R., Harun, M., Geraci, M., Carretero, M. A., Vinagre, C. and Tejedo, M. (2024). Does heat tolerance actually predict animals’ geographic thermal limits? Sci. Total Environ. 917, 170165.

Clark, R. M., Zera, A. J. and Behmer, S. T. (2014). Nutritional physiology of life history trade-offs: how food protein-carbohydrate content influences life-history traits in the wing-polymorphic cricket*Gryllus firmus*. J. Exp. Biol.

Clifford, C. W. and Woodring, J. P. (1990). Methods for rearing the house cricket, Acheta domesticus (L.), along with baseline values for feeding rates, growth rates, development times, and blood composition. J. Appl. Entomol. 109, 1–14.

Clifford, C. W., Roe, R. M. and Woodring, J. P. (1977). Rearing methods for obtaining house crickets, *Acheta domesticus*, of known age, sex, and instar. Ann. Entomol. Soc. Am. 70, 69–74.

Clissold, F. J. and Simpson, S. J. (2015). Temperature, food quality and life history traits of herbivorous insects. Curr. Opin. Insect Sci. 11, 63–70.

Deans, C. A. and Hutchison, W. (2023). The importance of time in nutrient regulation: a case study with spotted-wing Drosophila (Drosophila suzukii). *Front*. Insect Sci. 3, 1105531.

Deans, C. A., Sword, G. A. and Behmer, S. T. (2015). Revisiting macronutrient regulation in the polyphagous herbivore Helicoverpa zea (Lepidoptera: Noctuidae): New insights via nutritional geometry. J. Insect Physiol. 81, 21–27.

Deans, C. A., Sword, G. A., Vogel, H. and Behmer, S. T. (2022). Quantity versus quality: Effects of diet protein-carbohydrate ratios and amounts on insect herbivore gene expression. Insect Biochem. Mol. Biol. 145, 103773.

Desforges, J. E., BirnielJGauvin, K., Jutfelt, F., Gilmour, K. M., Eliason, E. J., Dressler, T. L., McKenzie, D. J., Bates, A. E., Lawrence, M. J., Fangue, N., et al. (2023). The ecological relevance of critical thermal maxima methodology for fishes. J. Fish Biol. 102, 1000–1016.

Dussutour, A. and Simpson, S. J. (2012). Ant workers die young and colonies collapse when fed a high-protein diet. Proc. R. Soc. B Biol. Sci. 279, 2402–2408.

Forster, J. and Hirst, A. G. (2012). The temperature-size rule emerges from ontogenetic differences between growth and development rates. Funct. Ecol. 26, 483–492.

Gibert, P. and De Jong, G. (2001). Temperature dependence of development rate and adult size in *Drosophila* species: biophysical parameters. J. Evol. Biol. 14, 267– 276.

González-Tokman, D., Córdoba-Aguilar, A., Dáttilo, W., Lira-Noriega, A., Sánchez-Guillén, R. A. and Villalobos, F. (2020). Insect responses to heat: physiological mechanisms, evolution and ecological implications in a warming world. Biol. Rev. 95, 802–821.

Grunert, L. W., Clarke, J. W., Ahuja, C., Eswaran, H. and Nijhout, H. F. (2015). A Quantitative Analysis of Growth and Size Regulation in Manduca sexta: The Physiological Basis of Variation in Size and Age at Metamorphosis. PLOS ONE 10, e0127988.

Hardison, E. A., Kraskura, K., Van Wert, J., Nguyen, T. and Eliason, E. J. (2021). Diet mediates thermal performance traits: implications for marine ectotherms. J. Exp. Biol. 224, jeb242846.

Jang, T. and Lee, K. P. (2018). Comparing the impacts of macronutrients on life-history traits in larval and adult Drosophila melanogaster: the use of nutritional geometry and chemically defined diets. J. Exp. Biol. 221, jeb181115.

Jensen, K., McClure, C., Priest, N. K. and Hunt, J. (2015). Sex-specific effects of protein and carbohydrate intake on reproduction but not lifespan in Drosophila melanogaster. Aging Cell 14, 605–615.

Johnson, J. A., Wofford, P. L. and Whitehand, L. C. (1992). Effect of Diet and Temperature on Development Rates, Survival, and Reproduction of the Indianmeal Moth (Lepidoptera: Pyralidae). J. Econ. Entomol. 85, 561–566.

Kaewtapee, C., Triwai, P., Inson, C., Masmeatathip, R. and Sriwongras, P. (2024). Effects of protein levels on production performance, nutritional values, and phase feeding of two-spotted cricket. J. Insect Sci. 24, 18.

Kim, K., Jang, T., Min, K. J. and Lee, K. P. (2020). Effects of dietary protein:carbohydrate balance on life history traits in six laboratory strains of Drosophila melanogaster. Entomol. Exp. Appl. 168, 482–491.

Kingsolver, J. G. and Pfennig, D. W. (2004). Individual-level selection as a cause of Cope’s rule of phyletic size increase. Evolution. 58, 1608–1612.

Kingsolver, J. G. and Umbanhowar, J. (2018). The analysis and interpretation of critical temperatures. J. Exp. Biol. 221, jeb.167858.

Kong, J. D., Vadboncoeur, É., Bertram, S. M. and MacMillan, H. A. (2025a). Temperature-dependence of life history in an edible cricket: Implications for optimising mass-rearing. Curr. Res. Insect Sci. 7, 100109.

Kong, J. D., Ritchie, M. W., Vadboncoeur, É., MacMillan, H. A. and Bertram, S. M. (2025b). Growth, development, and life history of a mass-reared edible insect, Gryllodes sigillatus (Orthoptera: Gryllidae). J. Econ. Entomol. 118, 1093–1103.

Lachenicht, M. W., Clusella-Trullas, S., Boardman, L., Le Roux, C. and Terblanche, J. S. (2010). Effects of acclimation temperature on thermal tolerance, locomotion performance and respiratory metabolism in Acheta domesticus L. (Orthoptera: Gryllidae). J. Insect Physiol. 56, 822–830.

Lamb, R. J. and Gerber, G. H. (1985). Effects of temperature on the development, growth, and survival of larvae and pupae of a north-temperate chrysomelid beetle. Oecologia 67, 8–18.

Lee, K. P. (2015). Dietary protein:carbohydrate balance is a critical modulator of lifespan and reproduction in *Drosophila melanogaster*: A test using a chemically defined diet. J. Insect Physiol. 75, 12–19.

Lee, K. P., Behmer, S. T. and Simpson, S. J. (2006). Nutrient regulation in relation to diet breadth: a comparison of Heliothis sister species and a hybrid. J. Exp. Biol. 209, 2076–2084.

Lee, K. P., Simpson, S. J. and Wilson, K. (2008a). Dietary protein-quality influences melanization and immune function in an insect. Funct. Ecol. 22, 1052–1061.

Lee, K. P., Simpson, S. J., Clissold, F. J., Brooks, R., Ballard, J. W. O., Taylor, P. W., Soran, N. and Raubenheimer, D. (2008b). Lifespan and reproduction in Drosophila: New insights from nutritional geometry. Proc. Natl. Acad. Sci. USA 105, 2498–2503.

Lee, K. P., Jang, T., Ravzanaadii, N. and Rho, M. S. (2015). Macronutrient balance modulates the temperature-size rule in an ectotherm. Am. Nat. 186, 212–222.

Leong, C.-M., Tsang, T. P. N. and Guénard, B. (2022). Testing the reliability and ecological implications of ramping rates in the measurement of critical thermal maximum. PLOS ONE 17, e0265361.

Li, C., Addeo, N. F., Rusch, T. W., Tarone, A. M. and Tomberlin, J. K. (2023). Black soldier fly (Diptera: Stratiomyidae) larval heat generation and management. Insect Sci. 30, 964–974.

Magara, H. J. O., Tanga, C. M., Fisher, B. L., Azrag, A. G. A., Niassy, S., Egonyu, J. P., Hugel, S., Roos, N., Ayieko, M. A., Sevgan, S., et al. (2024). Impact of temperature on the bionomics and geographical range margins of the two-spotted field cricket Gryllus bimaculatus in the world: Implications for its mass farming. PLOS ONE 19, e0300438.

Marshall, D. J., Pettersen, A. K., Bode, M. and White, C. R. (2020). Developmental cost theory predicts thermal environment and vulnerability to global warming. *Nat*. Ecol. Evol. 4, 406–411.

Mauritsson, K. and Jonsson, T. (2023). A new flexible model for maintenance and feeding expenses that improves description of individual growth in insects. Sci. Rep. 13, 16751.

Morales-Ramos, J. A., Macchiano, A. and Rojas, M. G. (2024). Estimating optimal temperature conditions for growth, development, and reproduction of Tenebrio molitor (Coleoptera: Tenebrionidae). J. Econ. Entomol. toae298.

Muzzatti, M. J., Harrison, S. J., McColville, E. R., Brittain, C. T., Brzezinski, H., Manivannan, S., Stabile, C. C., MacMillan, H. A. and Bertram, S. M. (2024). Applying nutritional ecology to optimize diets of crickets raised for food and feed. R. Soc. Open Sci. 11, 241710.

Nicholls, E., Rossi, M. and Niven, J. E. (2021). Larval nutrition impacts survival to adulthood, body size and the allometric scaling of metabolic rate in adult honeybees. J. Exp. Biol. 224, jeb242393.

Nijhout, H. F. and Callier, V. (2015). Developmental Mechanisms of Body Size and Wing-Body Scaling in Insects. Annu. Rev. Entomol. 60, 141–156.

Nufio, C. R., Sheffer, M. M., Smith, J. M., Troutman, M. T., Bawa, S. J., Taylor, E. D., Schoville, S. D., Williams, C. M. and Buckley, L. B. (2025). Insect size responses to climate change vary across elevations according to seasonal timing. PLOS Biol. 23, e3002805.

Nyamukondiwa, C., Weldon, C. W., Chown, S. L., le Roux, P. C. and Terblanche, J. S. (2013). Thermal biology, population fluctuations and implications of temperature extremes for the management of two globally significant insect pests. J. Insect Physiol. 59, 1199–1211.

Ørsted, M., Jørgensen, L. B. and Overgaard, J. (2022). Finding the right thermal limit: a framework to reconcile ecological, physiological and methodological aspects of CTmax in ectotherms. J. Exp. Biol. 225, jeb244514.

Overgaard, J., Kearney, M. R. and Hoffmann, A. A. (2014). Sensitivity to thermal extremes in Australian Drosophila implies similar impacts of climate change on the distribution of widespread and tropical species. Glob. Change Biol. 20, 1738–1750.

Peters, R. H. (1986). The Ecological Implications of Body Size. Cambridge, UK: Cambridge University Press.

Pinkert, S., Farwig, N., Kawahara, A. Y. and Jetz, W. (2025). Global hotspots of butterfly diversity are threatened in a warming world. *Nat*. Ecol. Evol. 9, 789– 800.

Quispe-Tarqui, R., Yujra Pari, J., Callizaya Condori, F. and Rebaudo, F. (2021). The Effect of Diet Interacting With Temperature on the Development Rate of a Noctuidae Quinoa Pest. Environ. Entomol. 50, 685–691.

Raubenheimer, D. and Simpson, S. J. (1999). Integrating nutrition: a geometrical approach. In Proceedings of the 10th International Symposium on Insect-Plant Relationships (eds Simpson, S. J., Mordue, A. J. and Hardie, J.), and Hardie, J.), pp. 67–82. Dordrecht: Springer Netherlands.

Raynal, R. S., Noble, D. W. A., Riley, J. L., Senior, A. M., Warner, D. A., While, G. M. and Schwanz, L. E. (2022). Impact of fluctuating developmental temperatures on phenotypic traits in reptiles: a meta-analysis. J. Exp. Biol. 225, jeb243369.

Rebaudo, F. and Rabhi, V.-B. (2018). Modeling temperature-dependent development rate and phenology in insects: review of major developments, challenges, and future directions. Entomol. Exp. Appl. 166, 607–617.

Régnière, J., Powell, J., Bentz, B. and Nealis, V. (2012). Effects of temperature on development, survival and reproduction of insects: Experimental design, data analysis and modeling. J. Insect Physiol. 58, 634–647.

Roe, R. M., Clifford, C. W. and Woodring, J. P. (1980). The effect of temperature on feeding, growth, and metabolism during the last larval stadium of the female house cricket, Acheta domesticus. J. Insect Physiol. 26, 639–644.

Roeder, K. A. and Behmer, S. T. (2014). Lifetime consequences of food protein-carbohydrate content for an insect herbivore. Funct. Ecol. 28, 1135–1143.

Shi, P. and Ge, F. (2010). A comparison of different thermal performance functions describing temperature-dependent development rates. J. Therm. Biol. 35, 225– 231.

Simpson, S. J. and Abisgold, J. D. (1985). Compensation by locusts for changes in dietary nutrients: behavioural mechanisms. Physiol. Entomol. 10, 443–452.

Simpson, S. J. and Raubenheimer, D. (1995). The geometric analysis of feeding and nutrition: a user’s guide. J. Insect Physiol. 41, 545–553.

Sissener, N. H., Hamre, K., Fjelldal, P. G., Philip, A. J. P., Espe, M., Miao, L., Høglund, E., Sørensen, C., Skjærven, K. H., Holen, E., et al. (2021). Can improved nutrition for Atlantic salmon in freshwater increase fish robustness, survival and growth after seawater transfer? Aquaculture 542, 736852.

Sripontan, Y., Chiu, C.-I., Tanansathaporn, S., Leasen, K. and Manlong, K. (2020). Modeling the Growth of Black Soldier Fly Hermetia illucens (Diptera: Stratiomyidae): An Approach to Evaluate Diet Quality. J. Econ. Entomol. 113, 742–751.

Thiex, N., Novotny, L. and Crawford, A. (2012). Determination of Ash in Animal Feed: AOAC Official Method 942.05 Revisited. J. AOAC Int. 95, 1392–1397.

Tjørve, K. M. C. and Tjørve, E. (2017). A proposed family of Unified models for sigmoidal growth. Ecol. Model. 359, 117–127.

Tu, X., Zhang, Z., Johnson, D. L., Cao, G., Li, Z., Gao, S., Nong, X. and Wang, G. (2012). Growth, Development and Daily Change in Body Weight of Locusta migratoria manilensis (Orthoptera: Acrididae) Nymphs at Different Temperatures. J. Orthoptera Res. 21, 133–140.

Weihrauch, D. and O’Donnell, M. J. (2021). Mechanisms of nitrogen excretion in insects. Curr. Opin. Insect Sci. 47, 25–30.

Whitman, D. W. (2008). The significance of body size in the Orthoptera: a review. J. Orthoptera Res. 17, 117–134.

Wojda, I. (2017). Temperature stress and insect immunity. J. Therm. Biol. 68, 96–103.

Zanotto, F. P., Simpson, S. J. and Raubenheimer, D. (1993). The regulation of growth by locusts through post-ingestive compensation for variation in the levels of dietary protein and carbohydrate. Physiol. Entomol. 18, 425–434.

Zanotto, F. P., Gouveia, S. M., Simpson, S. J., Raubenheimer, D. and Calder, P. C. (1997). Nutritional Homeostasis in LocUsts: Is There a Mechanism for Increased Energy Expenditure During Carbohydrate Overfeeding? J. Exp. Biol. 200, 2437– 2448.

Zera, A. J. and Harshman, L. G. (2001). The Physiology of Life History Trade-Offs in Animals. Annu. Rev. Ecol. Evol. Syst. 32, 95–126.

