## supplementary material, Table S1, Figure S1, Figure S2, Table S2 to S11 for "Temperature outweighs diet in shaping developmental performance in two cricket species via growth delays and physiological limits"

Supplemental material

Table S1. References supporting the conceptual framework in Figure 1, illustrating the typical response curves of insect life-history traits (survival, developmental rate, and adult mass) to variation in temperature and dietary protein-to-carbohydrate (P:C) ratio. Each reference is accompanied by a brief description of the relevant pattern it documents.

| **Reference** | **Summary Description** |
| --- | --- |
| Abarca et al., 2024 | Survival declines sharply near critical thermal limits in insects multiple *Lepidoptera* species. |
| Régnière et al., 2012 | Survival high across wide range of temperature and rapidly drops off at both ends of the range in *Choristoneura fumiferana* and *Bactrocera cucurbitae*. |
| Gibert and De Jong, 2001 | Developmental rate increases with temperature, peaks close to higher thermal limit and then either maintains or decreases slightly in four *Drosophila* species. |
| Lamb and Gerber, 1985 | Survival high across wide range of temperature and rapidly drops off at both ends of the range. Developmental rate increases with temperature, peaks close to higher thermal limit and is then maintained in *Entomoscelis americana*. |
| Morales-Ramos et al., 2024 | Developmental rate increases with temperature, peaks close to higher thermal limit and then decreases rapidly in *Tenebrio molitor*. |
| Atkinson, 1994 | Introduces the temperature-size rule in ectotherms. Most ectotherms, including many insect groups, will reach adulthood with larger mass at lower temperature. |
| Whitman, 2008 | Orthopterans (including crickets) deviate from the temperature-size rule and grow largest at intermediate to high temperature. |
| Kong et al., 2025 | Crickets grow largest at intermediate or high temperatures in *Gryllodes sigillatus*. |
| Dussutour and Simpson, 2012 | Survival decreases with high protein intake in *Lasius niger*. |
| Lee, 2015 | Survival decreases with high protein intake. Adult body weight increases with protein content in *Drosophila melanogaster*. |
| Lee et al., 2008 | High protein reduces survival in *Drosophila melanogaster*. |
| Bouchebti et al., 2022 | High protein reduces survival in *Blaptica dubia*. |
| Nicholls et al., 2021 | High protein reduces survival in *Apis mellifera*. |
| Kim et al., 2020 | Fastest development and adult mass on balanced to protein-biased diets. Lifespan maximized with low-protein diets in different strains of *Drosophila melanogaster*. |
| Muzzatti et al., 2024 | Developmental rate and mass at adulthood increased with protein content in feed while survival decreased in *Gryllodes sigillatus*. |
| Roeder and Behmer, 2014 | Development and mass maximized with moderate protein bias, while survival is maximized under carbohydrate-biased feeds in *Heliothis virescens*. |
| Kaewtapee et al., 2024 | Adult mass is highest under balanced P:C in *Gryllus bimaculatus*. |
| Jang & Lee 2018 | High developmental rate with protein-biased diets in *Drosophila melanogaster*. |

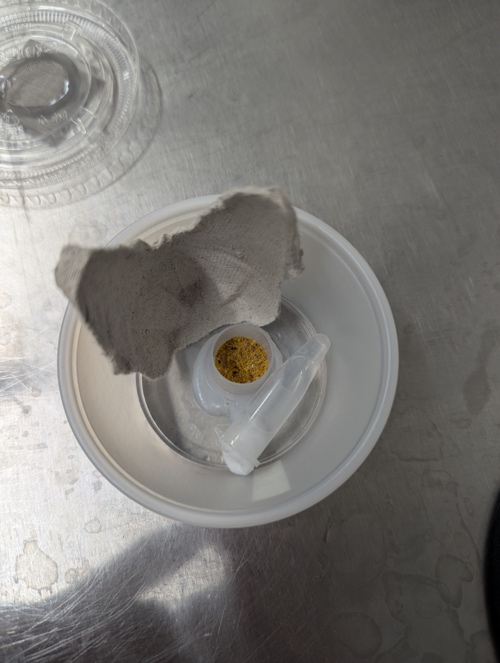

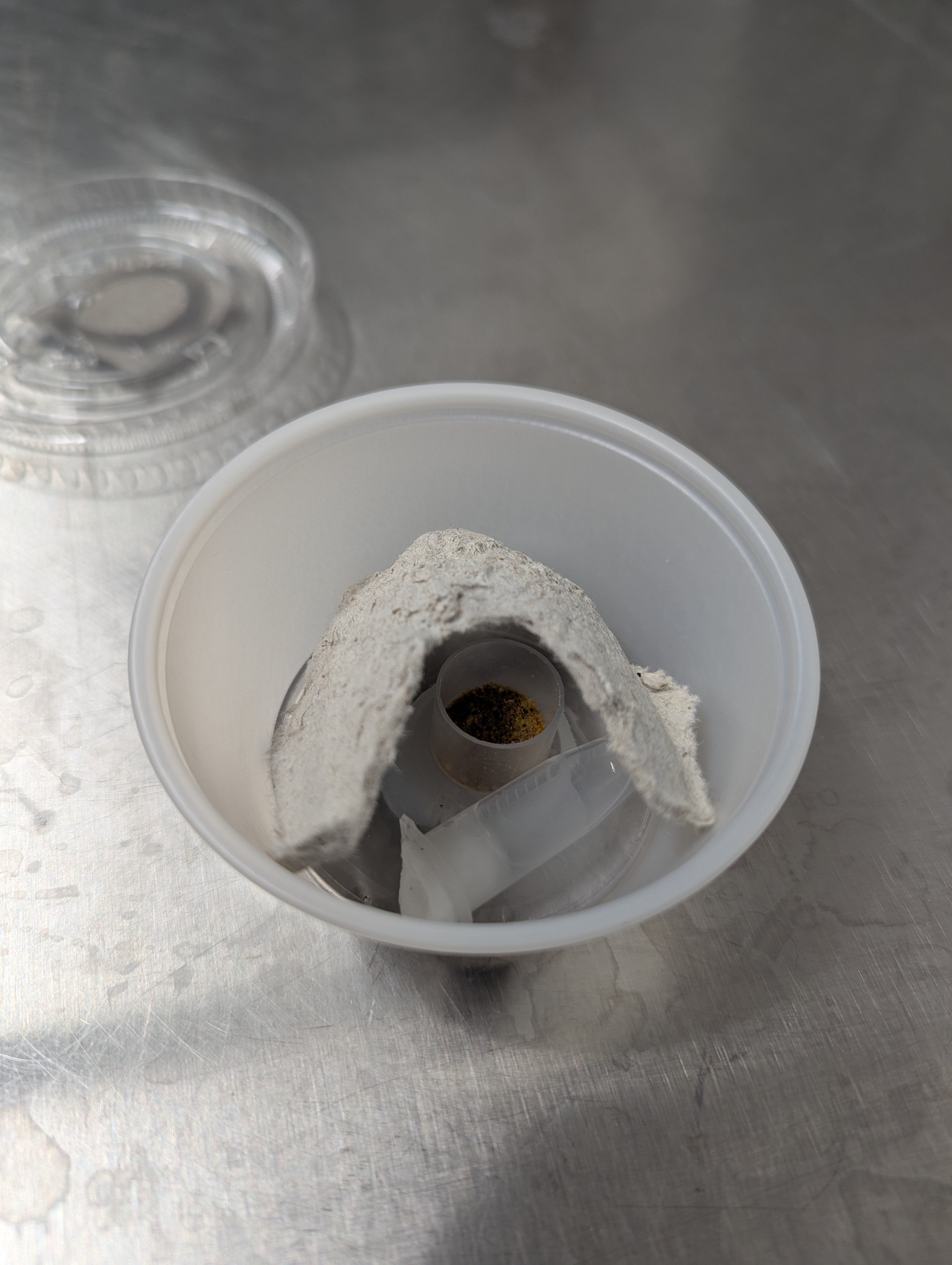

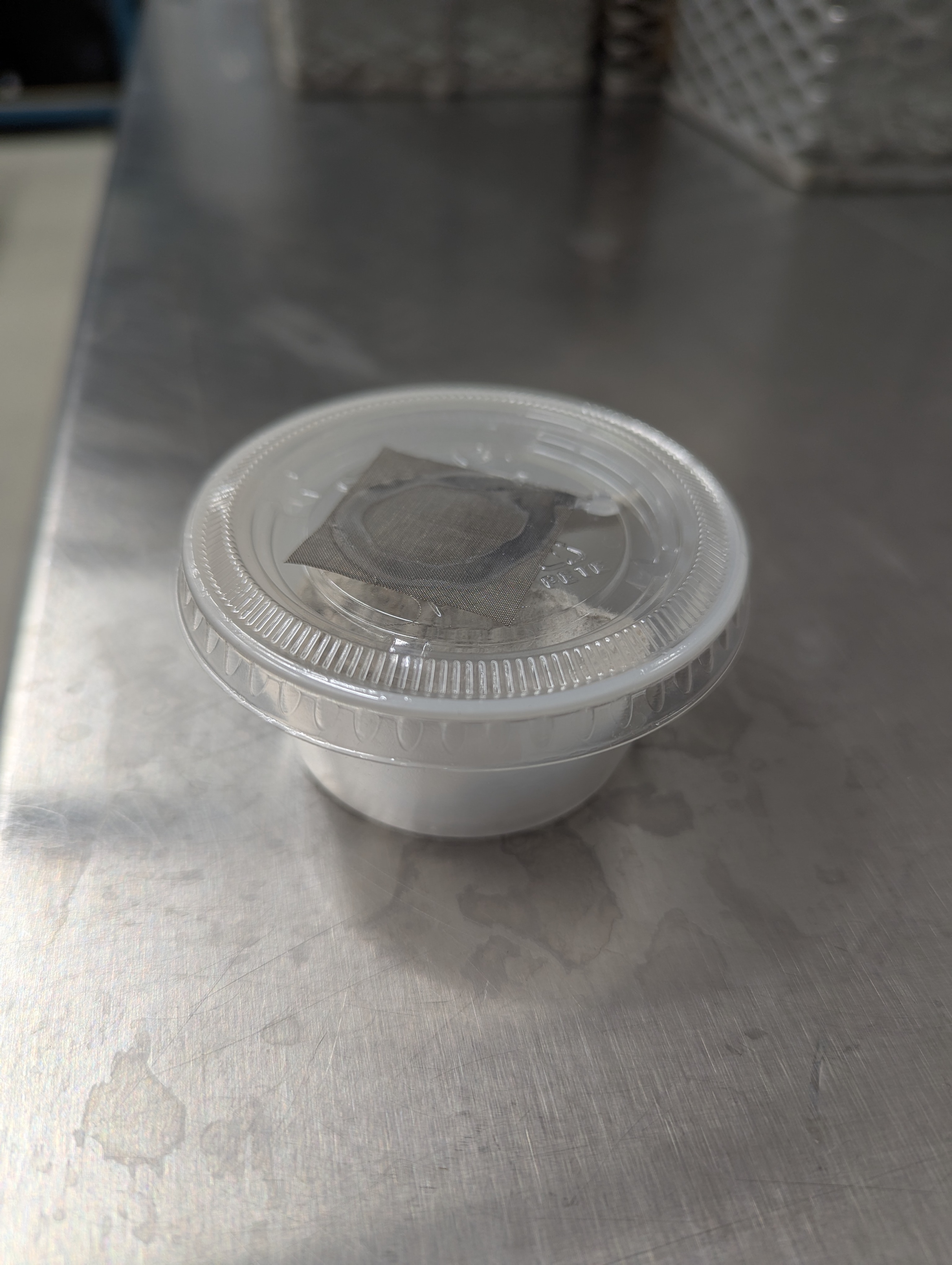

A

B

C

**Figure S1.** Rearing setup for individual crickets. (A) Closed 96 mL condiment container (7 cm diameter × 3 cm depth) with a ventilated plastic lid. (B) Interior view of the container showing a section of egg carton used as a shelter, a 1.5 mL microcentrifuge tube filled with water and sealed with dental cotton, and a small plastic ink cap (1.7 cm diameter × 1.4 cm depth) containing feed, fixed to a 50 mm Petri dish. (C) Shelter removed to show positioning of feed and water sources. Crickets were housed individually in these containers throughout the experiment.

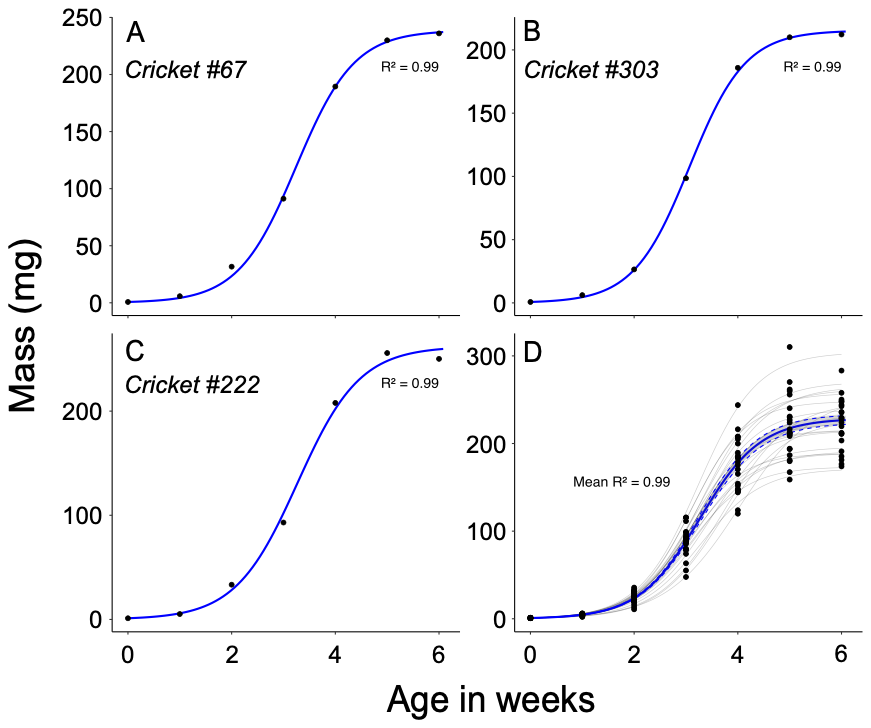

**Figure S2.** Representative examples of individual cricket growth modelled using **Equation 1**. Panels A–C show three randomly selected *G. sigillatus* males reared at 32 °C on a 0.83 protein-to-carbohydrate (P:C) diet, with body mass (mg) measured weekly over six weeks. Panel D shows growth trajectories for all individuals in this treatment group (black points), with fitted unified logistic curves overlaid (gray lines). Blue lines in panels A to C represent the model fit. Blue line in panel D represents a model made with the mean ± standard error of all crickets from the group. *R²* values are reported for each individual model, with a mean *R²* across the group, indicating strong model performance.

**Table S2.** Cox proportional hazards model output for *Acheta domesticus*, evaluating the effect of treatment on survival stratified by time group where strata1 represent day 0 to 17 and strata 2 day 18 to 42. Reported values include coefficient estimates (Coef.), hazard ratios, standard errors (SE), z values, and p-values (Pr(>|z|)).

| Predictors | Coef. | Hazard Ratio | Std. Error | z value | Pr(>\|z\|) |
| --- | --- | --- | --- | --- | --- |
| treatment4:strata1 | 1.144e+00 | 3.140e+00 | 1.118e+00 | 1.023 | 0.30617 |
| treatment1:strata1 | 1.444e+00 | 4.236e+00 | 1.118e+00 | 1.291 | 0.19662 |
| treatment2:strata1 | 2.069e+00 | 7.919e+00 | 1.069e+00 | 1.936 | 0.05291 . |
| treatment3:strata1 | 7.329e-01 | 2.081e+00 | 1.225e+00 | 0.598 | 0.54956 |
| treatment6:strata1 | 2.112e+00 | 8.263e+00 | 1.069e+00 | 1.975 | 0.04822 * |
| treatment8:strata1 | 1.439e+00 | 4.215e+00 | 1.118e+00 | 1.287 | 0.19816 |
| treatment9:strata1 | 2.448e+00 | 1.156e+01 | 1.049e+00 | 2.334 | 0.01962 * |
| treatment10:strata1 | 4.390e+00 | 8.066e+01 | 1.021e+00 | 4.299 | 1.718e-05 *** |
| treatment11:strata1 | -1.684e-02 | 9.833e-01 | 1.414e+00 | -0.012 | 0.99050 |
| treatment12:strata1 | NA | NA | 0.000e+00 | NA |  |
| treatment4:strata2 | -6.258e-09 | 1.000e+00 | 8.233e+03 | -0.000 | 1.00000 |
| treatment1:strata2 | -6.258e-09 | 1.000e+00 | 8.911e+03 | -0.000 | 1.00000 |
| treatment2:strata2 | -6.258e-09 | 1.000e+00 | 9.213e+03 | -0.000 | 1.00000 |
| treatment3:strata2 | 1.750e+01 | 3.990e+07 | 6.127e+03 | 0.003 | 0.99772 |
| treatment6:strata2 | -6.258e-09 | 1.000e+00 | 9.213e+03 | -0.000 | 1.00000 |
| treatment8:strata2 | 1.757e+01 | 4.286e+07 | 6.127e+03 | 0.003 | 0.99771 |
| treatment9:strata2 | 2.038e+01 | 7.114e+08 | 6.127e+03 | 0.003 | 0.99735 |
| treatment10:strata2 | NA | NA | 0.000e+00 | NA |  |
| treatment11:strata2 | -6.258e-09 | 1.000e+00 | 8.665e+03 | -0.000 | 1.00000 |
| treatment12:strata2 | NA | NA | 0.000e+00 | NA |  |
| Signif. codes: 0 ‘***’ 0.001 ‘**’ 0.01 ‘*’ 0.05 ‘.’ 0.1 ‘ ’ 1 | | | | | |

**Table S3.** Cox proportional hazards model output for *Gryllodes sigillatus*, evaluating the effect of treatment on survival stratified by time group where strata1 represent day 0 to 17 and strata 2 day 18 to 42. Reported values include coefficient estimates (Coef.), hazard ratios, standard errors (SE), z values, and p-values (Pr(>|z|)).

| Predictors | Coef. | Hazard Ratio | Std. Error | z value | Pr(>\|z\|) |
| --- | --- | --- | --- | --- | --- |
| treatment4:strata1 | -4.063e-01 | 6.661e-01 | 1.015e+00 | -0.400 | 0.68884 |
| treatment1:strata1 | 1.129e+00 | 3.094e+00 | 6.038e-01 | 1.870 | 0.06143 . |
| treatment2:strata1 | -1.226e+01 | 4.748e-06 | 4.554e+02 | -0.027 | 0.97852 |
| treatment3:strata1 | 7.101e-01 | 2.034e+00 | 6.038e-01 | 1.176 | 0.23958 |
| treatment5:strata1 | 7.303e-01 | 2.076e+00 | 7.282e-01 | 1.003 | 0.31596 |
| treatment6:strata1 | -1.226e+01 | 4.748e-06 | 4.554e+02 | -0.027 | 0.97852 |
| treatment7:strata1 | 1.898e+00 | 6.670e+00 | 4.485e-01 | 4.231 | 2.330e-05 *** |
| treatment8:strata1 | 7.303e-01 | 2.076e+00 | 7.282e-01 | 1.003 | 0.31596 |
| treatment9:strata1 | 6.920e-01 | 1.998e+00 | 7.282e-01 | 0.950 | 0.34202 |
| treatment10:strata1 | 2.389e+00 | 1.090e+01 | 3.743e-01 | 6.382 | 1.752e-10 *** |
| treatment11:strata1 | 1.727e+00 | 5.625e+00 | 4.831e-01 | 3.575 | 3.498e-04 *** |
| treatment12:strata1 | 0.000e+00 | 1.000e+00 | 1.015e+00 | 0.000 | 1.00000 |
| treatment4:strata2 | -1.742e+01 | 2.707e-08 | 4.854e+03 | -0.004 | 0.99714 |
| treatment1:strata2 | -1.742e+01 | 2.707e-08 | 6.197e+03 | -0.003 | 0.99776 |
| treatment2:strata2 | -1.742e+01 | 2.707e-08 | 5.879e+03 | -0.003 | 0.99764 |
| treatment3:strata2 | -3.958e-01 | 6.732e-01 | 1.026e+00 | -0.386 | 0.69957 |
| treatment5:strata2 | 3.756e-02 | 1.038e+00 | 1.026e+00 | 0.037 | 0.97079 |
| treatment6:strata2 | 6.495e-01 | 1.915e+00 | 7.446e-01 | 0.872 | 0.38304 |
| treatment7:strata2 | 8.866e-01 | 2.427e+00 | 7.444e-01 | 1.191 | 0.23365 |
| treatment8:strata2 | -1.742e+01 | 2.707e-08 | 6.085e+03 | -0.003 | 0.99772 |
| treatment9:strata2 | 5.872e-03 | 1.006e+00 | 1.026e+00 | 0.006 | 0.99543 |
| treatment10:strata2 | 3.362e+00 | 2.883e+01 | 4.539e-01 | 7.405 | 1.310e-13 *** |
| treatment11:strata2 | -1.742e+01 | 2.707e-08 | 6.440e+03 | -0.003 | 0.99784 |
| treatment12:strata2 | 0.000e+00 | 1.000e+00 | 1.026e+00 | 0.000 | 1.00000 |
| Signif. codes: 0 ‘***’ 0.001 ‘**’ 0.01 ‘*’ 0.05 ‘.’ 0.1 ‘ ’ 1 | | | | | |

**Table S4.** Generalized linear model output (Gamma distribution, log link) for *Acheta domesticus*, evaluating the effects of treatment and sex on asymptotic mass (A). Reported values include coefficient estimates, standard errors (SE), t values, and p-values (Pr(>|t|)).

| Predictors | Estimate | Std. Error | t value | Pr(>\|t\|) |
| --- | --- | --- | --- | --- |
| (Intercept) | 1.680e-03 | 6.757e-05 | 24.857 | < 2e-16 *** |
| treatment1 | 4.233e-04 | 1.313e-04 | 3.224 | 0.00145 ** |
| treatment2 | 7.767e-05 | 1.124e-04 | 0.691 | 0.49029 |
| treatment3 | 9.486e-05 | 1.091e-04 | 0.870 | 0.38537 |
| treatment6 | 2.980e-04 | 1.163e-04 | 2.563 | 0.01104 * |
| treatment8 | 7.462e-05 | 1.031e-04 | 0.724 | 0.46974 |
| treatment9 | 2.240e-03 | 2.108e-04 | 10.628 | < 2e-16 *** |
| treatment11 | 1.403e-04 | 1.004e-04 | 1.397 | 0.16367 |
| treatment12 | 5.042e-04 | 1.122e-04 | 4.494 | 1.124e-05 *** |
| sexfinalmale | 9.244e-04 | 6.795e-05 | 13.605 | < 2e-16 *** |
| Signif. codes: 0 ‘***’ 0.001 ‘**’ 0.01 ‘*’ 0.05 ‘.’ 0.1 ‘ ’ 1 | | | | |

**Table S5.** Generalized linear model output (Gamma distribution, log link) for *Gryllodes sigillatus*, evaluating the effects of treatment and sex on asymptotic mass (A). Reported values include coefficient estimates, standard errors (SE), t values, and p-values (Pr(>|t|)).

| Predictors | Estimate | Std. Error | t value | Pr(>\|t\|) |
| --- | --- | --- | --- | --- |
| (Intercept) | 2.832e-03 | 1.389e-04 | 20.395 | < 2e-16 *** |
| treatment1 | 3.146e-03 | 4.122e-04 | 7.631 | 2.592e-13 *** |
| treatment2 | 4.311e-04 | 2.229e-04 | 1.934 | 0.05400 . |
| treatment3 | 1.867e-04 | 1.968e-04 | 0.949 | 0.34337 |
| treatment5 | 1.544e-04 | 2.287e-04 | 0.675 | 0.50016 |
| treatment6 | 4.158e-04 | 2.365e-04 | 1.758 | 0.07968 . |
| treatment7 | 4.385e-04 | 2.566e-04 | 1.709 | 0.08846 . |
| treatment8 | -2.496e-04 | 2.020e-04 | -1.235 | 0.21755 |
| treatment9 | 3.535e-05 | 2.117e-04 | 0.167 | 0.86747 |
| treatment10 | 3.422e-03 | 4.887e-04 | 7.003 | 1.442e-11 *** |
| treatment11 | 4.458e-04 | 2.379e-04 | 1.874 | 0.06185 . |
| treatment12 | 1.094e-03 | 2.582e-04 | 4.235 | 2.978e-05 *** |
| sexfinalmale | 1.644e-03 | 1.176e-04 | 13.979 | < 2e-16 *** |
| Signif. codes: 0 ‘***’ 0.001 ‘**’ 0.01 ‘*’ 0.05 ‘.’ 0.1 ‘ ’ 1 | | | | |

**Table S6.** Generalized linear model output (Gamma distribution, log link) for *Acheta domesticus*, evaluating the effects of treatment, sex, and their interaction on the relative growth rate parameter (kU). Reported values include coefficient estimates, standard errors (SE), t values, and p-values (Pr(>|t|)).

| Predictors | Estimate | Std. Error | t value | Pr(>\|t\|) |
| --- | --- | --- | --- | --- |
| (Intercept) | 2.261e+00 | 3.757e-02 | 60.183 | < 2e-16 *** |
| treatment1 | 1.399e+00 | 9.384e-02 | 14.904 | < 2e-16 *** |
| treatment2 | 5.299e-01 | 7.558e-02 | 7.012 | 3.461e-11 *** |
| treatment3 | -1.793e-02 | 6.473e-02 | -0.277 | 0.78208 |
| treatment6 | 7.806e-02 | 5.918e-02 | 1.319 | 0.18866 |
| treatment8 | -2.809e-01 | 5.127e-02 | -5.479 | 1.265e-07 *** |
| treatment9 | -2.140e-01 | 5.911e-02 | -3.620 | 3.721e-04 *** |
| treatment11 | -1.073e-02 | 5.230e-02 | -0.205 | 0.83774 |
| treatment12 | 5.158e-02 | 5.301e-02 | 0.973 | 0.33170 |
| sexfinalmale | -2.138e-01 | 5.610e-02 | -3.811 | 1.840e-04 *** |
| treatment1:sexfinalmale | 2.031e-01 | 1.310e-01 | 1.550 | 0.12264 |
| treatment2:sexfinalmale | 2.675e-01 | 1.039e-01 | 2.574 | 0.01077 * |
| treatment3:sexfinalmale | 6.368e-02 | 8.489e-02 | 0.750 | 0.45400 |
| treatment6:sexfinalmale | 1.775e-01 | 8.873e-02 | 2.001 | 0.04675 * |
| treatment8:sexfinalmale | 1.621e-01 | 7.882e-02 | 2.057 | 0.04096 * |
| treatment9:sexfinalmale | 2.987e-01 | 8.797e-02 | 3.395 | 8.256e-04 *** |
| treatment11:sexfinalmale | -4.255e-03 | 8.389e-02 | -0.051 | 0.95959 |
| treatment12:sexfinalmale | 5.653e-02 | 8.279e-02 | 0.683 | 0.49554 |
| Signif. codes: 0 ‘***’ 0.001 ‘**’ 0.01 ‘*’ 0.05 ‘.’ 0.1 ‘ ’ 1 | | | | |

**Table S7.** Generalized linear model output (Gamma distribution, log link) for *Gryllodes sigillatus*, evaluating the effects of treatment, sex, and their interaction on the relative growth rate parameter (kU). Reported values include coefficient estimates, standard errors (SE), t values, and p-values (Pr(>|t|)).

| Predictors | Estimate | Std. Error | t value | Pr(>\|t\|) |
| --- | --- | --- | --- | --- |
| (Intercept) | 2.508e+00 | 5.167e-02 | 48.540 | < 2e-16 *** |
| treatment1 | 2.215e+00 | 1.724e-01 | 12.849 | < 2e-16 *** |
| treatment2 | 4.599e-01 | 8.407e-02 | 5.471 | 9.448e-08 *** |
| treatment3 | -7.856e-02 | 7.284e-02 | -1.078 | 0.28168 |
| treatment5 | 1.217e-01 | 8.947e-02 | 1.360 | 0.17470 |
| treatment6 | 1.020e-01 | 8.903e-02 | 1.146 | 0.25265 |
| treatment7 | 1.016e-01 | 9.947e-02 | 1.021 | 0.30795 |
| treatment8 | -3.330e-01 | 7.312e-02 | -4.554 | 7.638e-06 *** |
| treatment9 | -3.207e-01 | 7.216e-02 | -4.444 | 1.244e-05 *** |
| treatment10 | 1.413e-02 | 1.019e-01 | 0.139 | 0.88979 |
| treatment11 | 5.812e-02 | 8.162e-02 | 0.712 | 0.47698 |
| treatment12 | 5.036e-02 | 8.332e-02 | 0.604 | 0.54601 |
| sexfinalmale | -1.863e-01 | 6.891e-02 | -2.704 | 0.00724 ** |
| treatment1:sexfinalmale | -1.628e-01 | 2.104e-01 | -0.774 | 0.43973 |
| treatment2:sexfinalmale | 1.889e-02 | 1.233e-01 | 0.153 | 0.87833 |
| treatment3:sexfinalmale | 9.100e-02 | 9.791e-02 | 0.929 | 0.35345 |
| treatment5:sexfinalmale | -1.106e-01 | 1.147e-01 | -0.964 | 0.33605 |
| treatment6:sexfinalmale | 3.902e-03 | 1.138e-01 | 0.034 | 0.97267 |
| treatment7:sexfinalmale | 2.763e-01 | 1.280e-01 | 2.158 | 0.03169 * |
| treatment8:sexfinalmale | 9.599e-02 | 1.013e-01 | 0.948 | 0.34414 |
| treatment9:sexfinalmale | 1.865e-01 | 1.033e-01 | 1.805 | 0.07208 . |
| treatment10:sexfinalmale | 5.324e-01 | 1.498e-01 | 3.554 | 4.397e-04 *** |
| treatment11:sexfinalmale | -1.443e-01 | 1.160e-01 | -1.244 | 0.21439 |
| treatment12:sexfinalmale | 5.504e-02 | 1.134e-01 | 0.485 | 0.62788 |
| Signif. codes: 0 ‘***’ 0.001 ‘**’ 0.01 ‘*’ 0.05 ‘.’ 0.1 ‘ ’ 1 | | | | |

**Table S8.** Generalized linear model output (Gamma distribution, log link) for *Acheta domesticus*, evaluating the effects of treatment and sex on the slope of development. Reported values include coefficient estimates, standard errors (SE), t values, and p-values (Pr(>|t|)).

| Predictors | Estimate | Std. Error | t value | Pr(>\|t\|) |
| --- | --- | --- | --- | --- |
| (Intercept) | 6.265e-01 | 9.704e-03 | 64.564 | < 2e-16 *** |
| treatment1 | 3.872e-01 | 2.181e-02 | 17.756 | < 2e-16 *** |
| treatment2 | 1.945e-01 | 1.762e-02 | 11.043 | < 2e-16 *** |
| treatment3 | 1.633e-02 | 1.425e-02 | 1.146 | 0.25321 |
| treatment6 | 4.814e-02 | 1.535e-02 | 3.136 | 0.00195 ** |
| treatment8 | -8.177e-02 | 1.312e-02 | -6.231 | 2.320e-09 *** |
| treatment9 | 1.906e-02 | 1.622e-02 | 1.175 | 0.24106 |
| treatment11 | -3.118e-04 | 1.370e-02 | -0.023 | 0.98187 |
| treatment12 | 3.428e-02 | 1.411e-02 | 2.429 | 0.01594 * |
| sexfinalmale | 2.568e-02 | 7.852e-03 | 3.270 | 0.00125 ** |
| Signif. codes: 0 ‘***’ 0.001 ‘**’ 0.01 ‘*’ 0.05 ‘.’ 0.1 ‘ ’ 1 | | | | |

**Table S9.** Generalized linear model output (Gamma distribution, log link) for *Gryllodes sigillatus*, evaluating the effects of treatment, sex, and their interaction on the slope development. Reported values include coefficient estimates, standard errors (SE), t values, and p-values (Pr(>|t|)).

| Predictors | Estimate | Std. Error | t value | Pr(>\|t\|) |
| --- | --- | --- | --- | --- |
| (Intercept) | 6.733e-01 | 1.414e-02 | 47.610 | < 2e-16 *** |
| treatment1 | 5.086e-01 | 4.263e-02 | 11.929 | < 2e-16 *** |
| treatment2 | 1.013e-01 | 2.256e-02 | 4.491 | 1.005e-05 *** |
| treatment3 | -9.907e-03 | 1.985e-02 | -0.499 | 0.61811 |
| treatment5 | 3.806e-02 | 2.502e-02 | 1.521 | 0.12926 |
| treatment6 | 1.975e-02 | 2.459e-02 | 0.803 | 0.42245 |
| treatment7 | 2.990e-02 | 2.663e-02 | 1.123 | 0.26229 |
| treatment8 | -1.204e-01 | 1.972e-02 | -6.108 | 3.047e-09 *** |
| treatment9 | -8.558e-02 | 2.033e-02 | -4.210 | 3.364e-05 *** |
| treatment10 | 2.221e-02 | 3.077e-02 | 0.722 | 0.47089 |
| treatment11 | 6.952e-03 | 2.250e-02 | 0.309 | 0.75751 |
| treatment12 | 1.606e-02 | 2.221e-02 | 0.723 | 0.47020 |
| sexfinalmale | -2.401e-03 | 1.953e-02 | -0.123 | 0.90222 |
| treatment1:sexfinalmale | -9.289e-03 | 5.454e-02 | -0.170 | 0.86488 |
| treatment2:sexfinalmale | 1.916e-02 | 3.489e-02 | 0.549 | 0.58337 |
| treatment3:sexfinalmale | 6.432e-03 | 2.778e-02 | 0.232 | 0.81708 |
| treatment5:sexfinalmale | -5.803e-02 | 3.245e-02 | -1.789 | 0.07467 . |
| treatment6:sexfinalmale | 4.252e-03 | 3.216e-02 | 0.132 | 0.89491 |
| treatment7:sexfinalmale | 5.710e-02 | 3.605e-02 | 1.584 | 0.11426 |
| treatment8:sexfinalmale | 8.976e-03 | 2.849e-02 | 0.315 | 0.75291 |
| treatment9:sexfinalmale | 5.069e-02 | 3.113e-02 | 1.628 | 0.10455 |
| treatment10:sexfinalmale | 1.608e-01 | 4.747e-02 | 3.386 | 8.007e-04 *** |
| treatment11:sexfinalmale | 2.093e-03 | 3.281e-02 | 0.064 | 0.94918 |
| treatment12:sexfinalmale | 3.440e-02 | 3.233e-02 | 1.064 | 0.28820 |
| Signif. codes: 0 ‘***’ 0.001 ‘**’ 0.01 ‘*’ 0.05 ‘.’ 0.1 ‘ ’ 1 | | | | |

**Table S10.** Generalized linear model output (Gamma distribution, log link) for *Acheta domesticus*, evaluating the effect of treatment on developmental performance. Reported values include coefficient estimates, standard errors (SE), t values, and p-values (Pr(>|t|)).

| Predictors | Estimate | Std. Error | t value | Pr(>\|t\|) |
| --- | --- | --- | --- | --- |
| (Intercept) | 1.142e+00 | 4.518e-02 | 25.281 | < 2e-16 *** |
| treatment1 | 9.652e-01 | 1.192e-01 | 8.100 | 3.644e-14 *** |
| treatment2 | 5.611e-01 | 9.459e-02 | 5.932 | 1.139e-08 *** |
| treatment3 | -4.390e-03 | 6.762e-02 | -0.065 | 0.94829 |
| treatment6 | 4.361e-01 | 8.928e-02 | 4.885 | 1.983e-06 *** |
| treatment8 | -4.455e-02 | 6.766e-02 | -0.658 | 0.51094 |
| treatment9 | 4.864e+00 | 3.343e-01 | 14.548 | < 2e-16 *** |
| treatment11 | -4.134e-02 | 6.579e-02 | -0.628 | 0.53039 |
| treatment12 | 2.137e-01 | 7.424e-02 | 2.878 | 0.00439 ** |
| Signif. codes: 0 ‘***’ 0.001 ‘**’ 0.01 ‘*’ 0.05 ‘.’ 0.1 ‘ ’ 1 | | | | |

**Table S11.** Generalized linear model output (Gamma distribution, log link) for *Gryllodes sigillatus*, evaluating the effect of treatment on developmental performance. Reported values include coefficient estimates, standard errors (SE), t values, and p-values (Pr(>|t|)).

| Predictors | Estimate | Std. Error | t value | Pr(>\|t\|) |
| --- | --- | --- | --- | --- |
| (Intercept) | 1.022e+00 | 4.152e-02 | 24.609 | < 2e-16 *** |
| treatment1 | 2.696e+00 | 2.226e-01 | 12.112 | < 2e-16 *** |
| treatment2 | 2.971e-01 | 7.798e-02 | 3.810 | 1.670e-04 *** |
| treatment3 | 1.170e-01 | 6.299e-02 | 1.858 | 0.06409 . |
| treatment5 | 1.311e-01 | 7.280e-02 | 1.801 | 0.07272 . |
| treatment6 | 2.183e-01 | 7.468e-02 | 2.923 | 0.00371 ** |
| treatment7 | 6.612e-01 | 1.052e-01 | 6.282 | 1.095e-09 *** |
| treatment8 | -2.257e-01 | 5.856e-02 | -3.854 | 1.404e-04 *** |
| treatment9 | -3.015e-02 | 6.769e-02 | -0.445 | 0.65635 |
| treatment10 | 7.728e+00 | 6.821e-01 | 11.330 | < 2e-16 *** |
| treatment11 | 3.553e-01 | 8.506e-02 | 4.177 | 3.809e-05 *** |
| treatment12 | 4.529e-01 | 8.583e-02 | 5.277 | 2.429e-07 *** |
| Signif. codes: 0 ‘***’ 0.001 ‘**’ 0.01 ‘*’ 0.05 ‘.’ 0.1 ‘ ’ 1 | | | | |
